# Transfer potential of F-like plasmids in *Escherichia coli* differs by animal environment

**DOI:** 10.64898/2026.02.02.703319

**Authors:** Sneha Sundar, Sebastian Bonhoeffer, Jana S. Huisman

## Abstract

Plasmids play a key role in the spread of virulence and antimicrobial resistance genes to new genetic backgrounds. Genetic variation in the transfer operon, which contain the genes responsible for conjugation, can lead to substantial differences in transfer potential even between closely related plasmids. However, it is not clear how much genetic diversity there is in transfer operons of natural bacterial populations. Here, we analyse the prevalence and transfer potential of F-like plasmids, a clinically important family of plasmids in *Enterobacteriaceae*, by searching for the presence of essential conjugation genes in 1200 *Escherichia coli* genomes isolated from three livestock-associated environments. We find that the fraction of F-like transfer operons that are functionally complete was significantly higher in poultry than in bovine and swine associated bacteria. This difference was not captured in methods that use the presence of replication genes to estimate plasmid prevalence. Confounders such as the phylogenetic relatedness of *E. coli* or the presence of antibiotic resistance could not explain these significant differences in transfer potential. Instead, it seems the poultry environment selects for plasmids with high transfer potential, as it also contained more conjugative plasmid types per isolate. While we find environment specific differences in overall plasmid frequency, patterns of transfer gene presence/absence were similar across the three environments. Regulatory and exclusion genes are the exception to this pattern, suggesting environment specific modulation of transfer rates. This highlights the use of genomic data to uncover environment specific differences in plasmid prevalence and transfer potential, revealing the selection pressures shaping horizontal gene transfer in these environments.

## 1 Introduction

Plasmids are key vectors in the spread of clinically-important traits such as antibiotic resistance, virulence and metabolic genes to new genetic backgrounds [1–5]. A plasmid’s ability to transfer is contingent on the presence of functional transfer genes (*tra* genes), which encode and regulate the structures necessary for conjugation [6]. For a few model plasmids the mechanistic basis of conjugation has been studied extensively *in vitro* [7, 8], yet little is known about the genetic diversity and function of transfer genes in natural environments. This limits our ability to predict how plasmids spread in natural bacterial populations, and whether there are any differences depending on the ecological context.

The genetic diversity of plasmids is shaped by environment-specific evolutionary selection pressures. Specifically, a plasmid must balance the growth cost it places on the bacterial host with horizontal spread and selection for plasmid-borne genes to optimize their reproductive fitness in a given environment [9–11]. Conjugative plasmid carriage and the process of conjugation impose costs on the host cell [10, 12]. In experimental settings, these costs are compensated over time by bacterial host or plasmid mutations [12–14] or even the loss of transfer genes [15]. Phages that target conjugative pili for entry also select for plasmids that lack functional conjugation machinery [16] and such phages are abundant in natural samples [17].

These selection pressures may alter a plasmid’s ability to transfer, i.e. its transfer potential. Experiments have shown that a loss of structural genes involved in forming the type IV secretion system (T4SS) renders a plasmid incapable of conjugating on its own [7, 8]. Loss of regulatory genes has been reported to affect conjugation efficiency [18, 19], while loss of exclusion genes may increase super infection of closely-related plasmids in the same host cell [20, 21]. Depending on the genes that are lost, the plasmid may still be able to transfer if another plasmid with a functional conjugation system is present (termed mobilizable plasmid), or be rendered completely immobile (termed non-mobilizable plasmid). In natural samples, mobilizable plasmids are common, representing 21% of all plasmids [22].

Plasmids are thought to transition from conjugative to mobilizable states frequently [22, 23], and a diversity of transfer operon structures exists among closely-related plasmids with plasmids often missing one or more transfer genes [24, 25]. Since the genomic presence/absence of *tra* genes has been shown to be a good determinant of transfer potential in experiments with natural IncF and IncI plasmids [26], these genes may serve as useful functional markers to estimate both plasmid prevalence and transfer potential.

Existing studies that investigate natural plasmid diversity in relation to mobility usually suffer from one of three issues. First, they rely on sequence databases which may represent a biased sample of plasmid diversity: clinically relevant and experimentally used plasmids tend to be overrepresented, and the plasmids do not necessarily represent the plasmid populations found in a well-defined ecological niche [22, 24, 27–29]. Second, plasmid prevalence is often assessed based solely on the replicon sequence [30–33]. While the replicon is useful to define incompatibility groups, it is less informative about transfer potential. Third, methods designed to assess plasmid mobility typically use very relaxed thresholds to assess transfer potential. Existing mobility typing schemes have either used (i) just the relaxase, which only distinguishes mobilizable plasmids from non-mobilizable ones [34, 35], or (ii) the relaxase and a small number of other conjugation-related genes [22, 36, 37]. The latter approach can distinguish conjugative, mobilizable and non-mobilizable plasmids. Yet in practice, the thresholds used to classify a plasmid as conjugative are not specific to the conjugation system assessed and are lacking a functional basis. For instance, bioinformatic studies on plasmid mobility have used as little as three mating pair formation system (MPF) genes to call F-like plasmids conjugative [22, 36], while experiments have demonstrated that more than ten MPF genes are required for conjugation [6–8].

Here, we study the prevalence, genetic diversity and transfer potential of F-like plasmids in natural *Escherichia coli*(*E. coli*) populations. We analyze a collection of 1200 *E. coli* genomes in three livestock-associated environments (swine, poultry and cattle) for the presence of 35 F-like transfer genes (Figure 1A). We define a new metric of transfer operon completeness which is functionally linked to transfer potential, and find notable differences in plasmid prevalence and completeness across the three livestock-associated environments. This has important epidemiological implications for the spread of plasmid-borne resistance in *Enterobacteriaceae*. Our work also highlights the benefits of integrating molecular insights into bioinformatic analyses, to elucidate evolutionary selection pressures influencing plasmid spread in natural environments.

**Figure 1:**
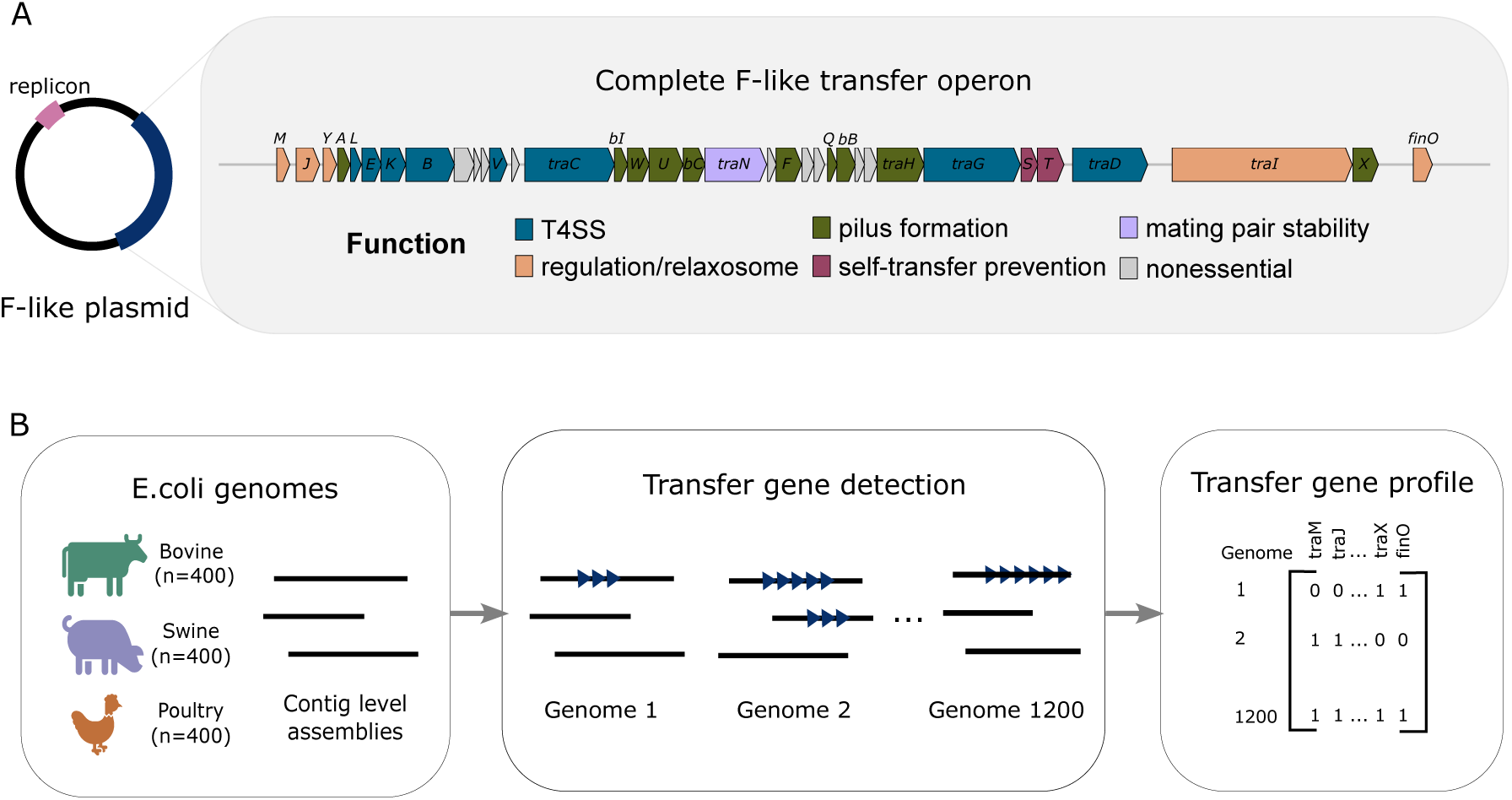
**A.** Schematic of the F-like plasmid with a replicon (pink) and an F-like transfer operon (dark blue). A magnified view of the operon displays individual genes, where colours indicate the function of different genes in the operon. **B.** Bioinformatic pipeline to detect transfer genes: (i) we collected 1200 contig-level genome assemblies associated with three animal environments, (ii) used a combined sequence homology and annotation based approach to identify a set of 35 genes from the F plasmid transfer operon, and (iii) determined a presence/absence profile of F-like transfer genes for each genome.

## 2 Material and Methods

### 2.1 *E. coli* Dataset

We used a collection of 1200 publicly available *E. coli* genomes associated with livestock, sampled as part of routine surveillance from USA between years 2010 and 2018 (Figure S1). The genomes used were not preferentially isolated based on virulence, resistance or strain characteristics. Our collection represents a subset of the dataset used in [38], randomly subsampled to select equal numbers (n=400 each) of bovine, swine and poultry-associated genomes, corresponding to the number of swine-associated samples available. The genomes had all been processed with a common assembly pipeline from Enterobase to generate contig-level assemblies and passed Enterobase’s internal assembly quality standards [39, 40], which include *E. coli*-specific criteria for the number of bases in the assembly, N50 value, number of contigs, proportion of scaffolding placeholders and species assignment [40]. The quality of the draft genomes were also independently assessed using checkM2 (v.1.1.0, using default settings) [41] and were found to be high quality (≥ 99.9% completeness, *<*3% contamination). Details on the metadata associated with the genomes — phylogroup, antimicrobial resistance gene (ARG) presence, sample source and sampling year — can be found in the Supplementary Text.

### 2.2 Detection of F-like transfer genes from short-read bacterial assemblies

To identify F-like transfer genes, we first used a sequence similarity based gene finding approach using a reference database of transfer gene sequences from *Enterobacteriaceae* F-like plasmids adapted from Fernandez-Lopez *et al* [24] (see Supplementary Text). This reference dataset was comprised of sequence homologues of 35 F-like *tra* genes (Table S1).

Using this set of *tra* homologues as queries, we ran tblastn (BLAST+ 2.12.0) [42] to identify F-like transfer operon associated contigs in the *E. coli* genomes. To allow for distant homologues, we selected hits with a minimum percent identity (pid) of 50%. To allow partial genes or genes on the edge of contigs to be detected, we selected a minimum query coverage of 30%. The best hit for each of the 35 *tra* marker genes (defined as the hit with the highest bitscore, pid and query coverage, prioritized in this order) was selected from each contig. In this way, contigs containing one or more F-like transfer genes were identified as likely F-like plasmid-associated contigs. We found an excess of contigs with only the transfer gene *traT*. As chromosomal homologues of *traT* have been reported before [43], we inspected the other genes on these contigs and found them to be chromosome-associated. We filtered out such contigs with *traT* as the only *tra* gene.

We then used the bacterial genome and plasmid annotation tool Bakta (v. 1.8.1, with default settings) [44] to independently cross-check the hits found by our *tra* gene-specific BLAST. We used the annotations shown in Table S1 to identify F-like transfer genes. Overall, we found little difference between the Bakta and BLAST approach in *tra* gene presence/absence. However, Bakta was better in identifying the entry exclusion gene *traS*, which has been described to have low sequence homology [24, 45]. For this reason, we used the set of transfer genes identified by Bakta in subsequent analysis.

### 2.3 Measuring prevalence of F-like transfer operons, replicons and transfer genes

An F-like transfer operon was determined as present if more than a given number of F-like transfer genes (the detection threshold) were detected in the genome. Prevalence of F-like transfer operons, defined as the ratio of the number of genomes containing an F-like transfer operon to the total number of genomes assessed, was evaluated for a range of detection thresholds (1-35 genes).

Replicon sequences in each genome were detected using the tool PlasmidFinder (v. 2.1.6) [46] with default parameters (minimum identity 95% and minimum coverage 60%). PlasmidFinder detects plasmids in genomes using a pre-defined database of replicon sequences as queries in a BLAST search. The genome was said to contain an F-like replicon if one or more replicons of type IncFIA, IncFIB, IncFIC or IncFII were detected. Prevalence of F-like replicons was defined as the number of genomes containing an F-like replicon divided by the total number of genomes assessed.

We estimated transfer gene frequencies using a gene content matrix with 1200 isolates (rows) and 35 transfer genes (columns), where each entry represents the number of detected hits for a given gene in an isolate. A binary presence–absence matrix was derived from this matrix (1 = present, 0 = absent). Gene frequencies were calculated as the proportion of genomes in which each gene was detected, conditioned on genomes containing at least one transfer gene (n=249 in bovine, n=350 in poultry, n=273 in swine) to control for differences in the proportion of genomes lacking transfer genes across animal sources.

### 2.4 Classifying transfer potential of F-like transfer operons

We used molecular insights from experimental studies to classify the transfer potential of F-like plasmids based on the presence/absence of a set of 23 essential transfer genes (Table S1 and Supplementary Text).

Every genome was classified as containing a ”complete”, ”incomplete” or ”missing” F-like transfer operon depending on the number of essential genes detected in the genome. Genomes were classified as ”missing” F-like transfer operons if *<* 5 genes from the essential gene set were detected, as ”incomplete” if *>*= 5 but *<* 23 genes were detected, and ”complete” if all 23 essential genes were detected.

### 2.5 Logistic regression model

The presence of a complete F-like operon was modelled using logistic regression implemented as a generalized linear model with a binomial error distribution using the function *glm* in R version 4.2.2. Candidate explanatory variables included Animal Environment (poultry, bovine, swine), Phylogroup (A–F), Sample Source (food, livestock), antibiotic resistance gene (ARG) presence (yes,no), and sampling year (before 2017, 2017 or later).

Model selection was performed using Akaike’s Information Criterion (AIC). All combinations of explanatory variables were tested to identify the main-effects model with the lowest AIC (Table S4). Pairwise interaction terms were then added sequentially to the selected main-effects model, and again the model with the lowest AIC was selected (Table S5).

The final selected model included Animal Environment, Phylogroup, Sample Source, and ARG presence as main effects, as well as an interaction between Phylogroup and ARG presence. Pairwise contrasts were computed using the *emmeans* package (v. 1.10.5) [47] with this selected model and multiple testing correction was performed using the Tukey method.

### 2.6 Detecting non-F conjugation systems

We used geNOMAD (v. 1.7.4) [48] to identify the presence of non-F conjugation systems in our isolates. We computed two binary matrices (1 = present, 0 = absent) for mating pair formation system (MPF) type presence (1200 x 4) and relaxase type presence (1200 x 6). An MPF type is considered present if we could detect at least 3 distinct genes specific to that MPF type. This is analogous to how we defined F-like transfer operon presence but scaled to accommodate transfer systems with fewer MPF genes. A relaxase type is termed present in an isolate if we could detect at least one gene of that type in the isolate. Using these matrices, we computed the number of non-F MPF types per isolate and non-F relaxase types per isolate. The prevalence of each MPF type and relaxase type was also calculated from this binary matrix as the proportion of genomes in which an MPF or relaxase type was detected.

### 2.7 Data Availability

Code and data files (including the accession numbers of genomes) used to reproduce the analyses and figures can be found in this github repository https://github.com/ Sneha-Sundar/F_div_paper.

## 3 Results

### 3.1 F-like transfer operons are less prevalent than IncF replicons in *E. coli* from livestock

We set out to study the prevalence and genetic diversity of F-like plasmids in natural environments. To do so, we used a collection of 1200 publicly available *E. coli* genomes from the USA associated with three livestock environments [38]: bovine (*n* = 400), swine (*n* = 400) and poultry (*n* = 400). The isolates were sampled as part of routine farm surveillance by the USA Food and Drug Administration between 2010 and 2018 (Figure S1) and were not specifically isolated based on any virulence, strain or resistance characteristics. On top of passing assembly quality standards set by Enterobase [40], the quality of the draft genomes were independently assessed using checkM2 [41] and were found to be high quality (Figure S2). All 1200 genomes had a completeness score ≥ 99.9%, out of which 1194 genomes were complete (100% completeness score).

We assessed how similar the genomes were in terms of their pairwise average nucleotide identity (ANI). The ANI distribution follows the typical patterns expected for *E. coli* [49]: all pairs had ANI *>* 96%, genomes belonging to the same PhG had average pairwise ANI of 99%, and the same ST 99.5% (Figures S3A,B). We found 42 clusters of highly similar genomes (121 genomes in total or 10% of the dataset) with ANI values ≥ 99.99%, which meets a recommended ANI threshold for strain identification [50] (Figure S3C). However, these potential strains could still have substantial differences in mobile genetic element (MGE) content [49, 51], which are not captured using ANI. Highly similar genomes may also be sampled because they are more prevalent. Therefore, for most analyses presented we do not perform any genome dereplication.

We developed a method using a combined sequence homology and annotation based approach to detect a set of 35 genes from the F plasmid transfer operon in our *E. coli* genomes (Figure 1). This resulted in a presence/absence profile of F-like transfer genes for each genome (Figure 1B). Since we work with contig-level assemblies, plasmid sequences are likely to be fragmented across multiple contigs (Figure S4 [52, 53]. We bypass this structural limitation by using a gene-based detection method, which successfully identifies F-like transfer operons regardless of contig fragmentation.

In the following, we use the F-like transfer operon both as a marker to detect the presence of the F-like plasmid and to assess transfer potential.

We first set out to estimate the prevalence of F-like plasmids in our *E. coli* genomes. Using a minimum threshold of 5 F-like transfer genes to mark an F-like plasmid as present, we found a prevalence of 65.5% in the entire dataset. We found significant differences in F-like plasmid prevalence between the three animal environments (χ^2^(2) = 152.00*,p <* 2*e* — 16; Figure 2A) with bovine at 44.75%. swine at 65.5%, and poultry at 86.25%. These prevalence estimates were robust to a wide range of marker gene thresholds (Figure 2B). At the low end (*<* 4) there is a sub-population of bovine *E. coli* with 3 transfer genes that inflate the number of F-like transfer operons identified. On the high end, prevalence begins to drop quickly above 27 marker genes in all three animal environments. Interestingly, this number is larger than the 23 genes with a demonstrated function in F conjugation [3, 7, 8]. Only 2.67% of all genomes contained all 35 marker genes found in the classical F plasmid.

**Figure 2:**
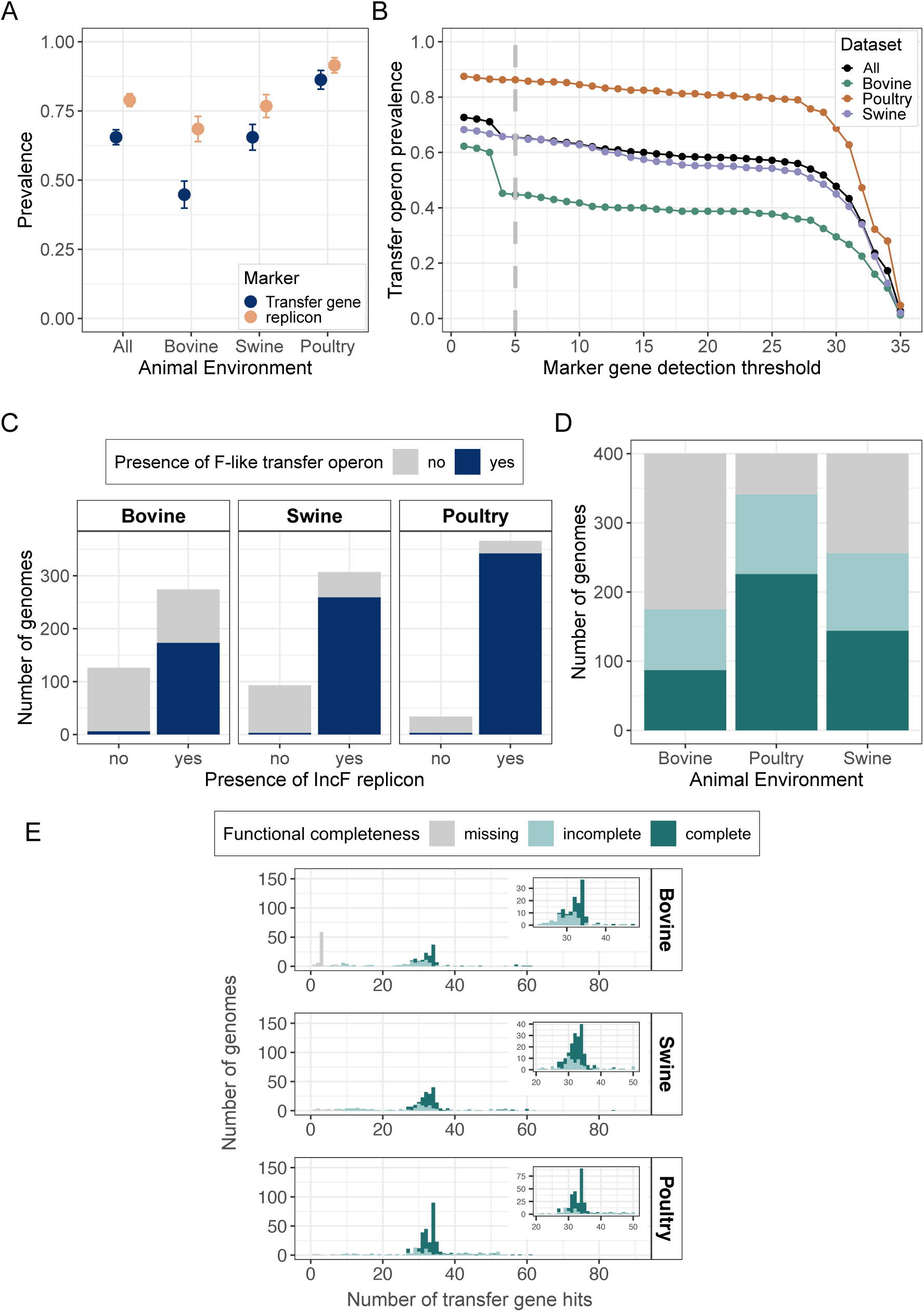
**A.** Prevalence of F-like plasmids estimated using transfer genes (blue) or replicons (orange) as markers across the entire dataset (all; n = 1200 *E. coli* isolates) and within each animal environment (n = 400 each). A minimum of 5 different transfer genes were necessary to call a plasmid ‘present’ in a genome. **B.** Estimates of F-like transfer operon prevalence across our entire dataset (dark blue) and in each animal environment (Bovine: green, Poultry: orange, Swine: purple) as a function of the number of genes required to mark an operon as present. The vertical dashed line indicates the 5 gene detection threshold used in further analyses. **C.** Proportion of *E. coli* genomes containing F-like transfer operons (blue) or not (grey), given the presence or absence of an IncF replicon in a genome. **D.** Proportion of *E. coli* isolates from each animal environment that have F-like transfer operons that are ‘complete’ (all 23 essential F-like transfer genes), ‘incomplete’ (between 5 and 22 essential genes) or ‘missing’ (less than 5 essential genes). **E.** Number of F-like transfer genes detected per genome across the three animal environments. Genomes are colored by the functional completeness category they were assigned. The insets show a zoom-in of the distribution between 20 and 50 transfer gene hits. Colour legends for panels D and E are the same.

To compare to past estimates of plasmid prevalence in environmental datasets, we also estimated the prevalence of IncF replicons in our dataset (2A). Replicons are loci involved in replication (Figure 1A), and are very commonly used to classify plasmids into families that reflect incompatibility groupings. Using the replicon typing tool Plas-midFinder, IncF replicon prevalence was found to be 81.22% in the entire dataset and 68.5%, 76.75%, and 91.50% in bovine, swine and poultry respectively. This difference was statistically significant between animal environments: χ^2^(2) = 65*,p <* 7*e* — 15. The replicon-based prevalence estimates were always higher than those based on the F-like transfer operon (Figure 2A). Absence of a replicon was associated with absence of the transfer operon(Figure 2C)). The inverse, however, was not necessarily true (Figure 2C) as 36.86%, 15.64%, 6.56% of bovine, swine and poultry *E. coli* that contain IncF replicons were lacking F-like transfer operons.

Our estimate of replicon-based prevalence of 81.22% is similar to previously reported prevalence estimates in livestock and human commensals (85.2% and 82.19% respectively [31, 32]; Table 1), and higher than estimates reported for wastewater treatment plants and human bloodstream infections (73.5% and 73.51% respectively [31, 33]). However, our transfer gene based prevalence estimate (65.5%) is lower than replicon based prevalence estimates reported across all datasets. Interestingly, both our transfer-gene and replicon-based prevalence estimates vary across the three animal environments, in contrast to a livestock dataset from England [31], which reported similar replicon-based prevalences across the same livestock environments (Table 1).

**Table 1:**
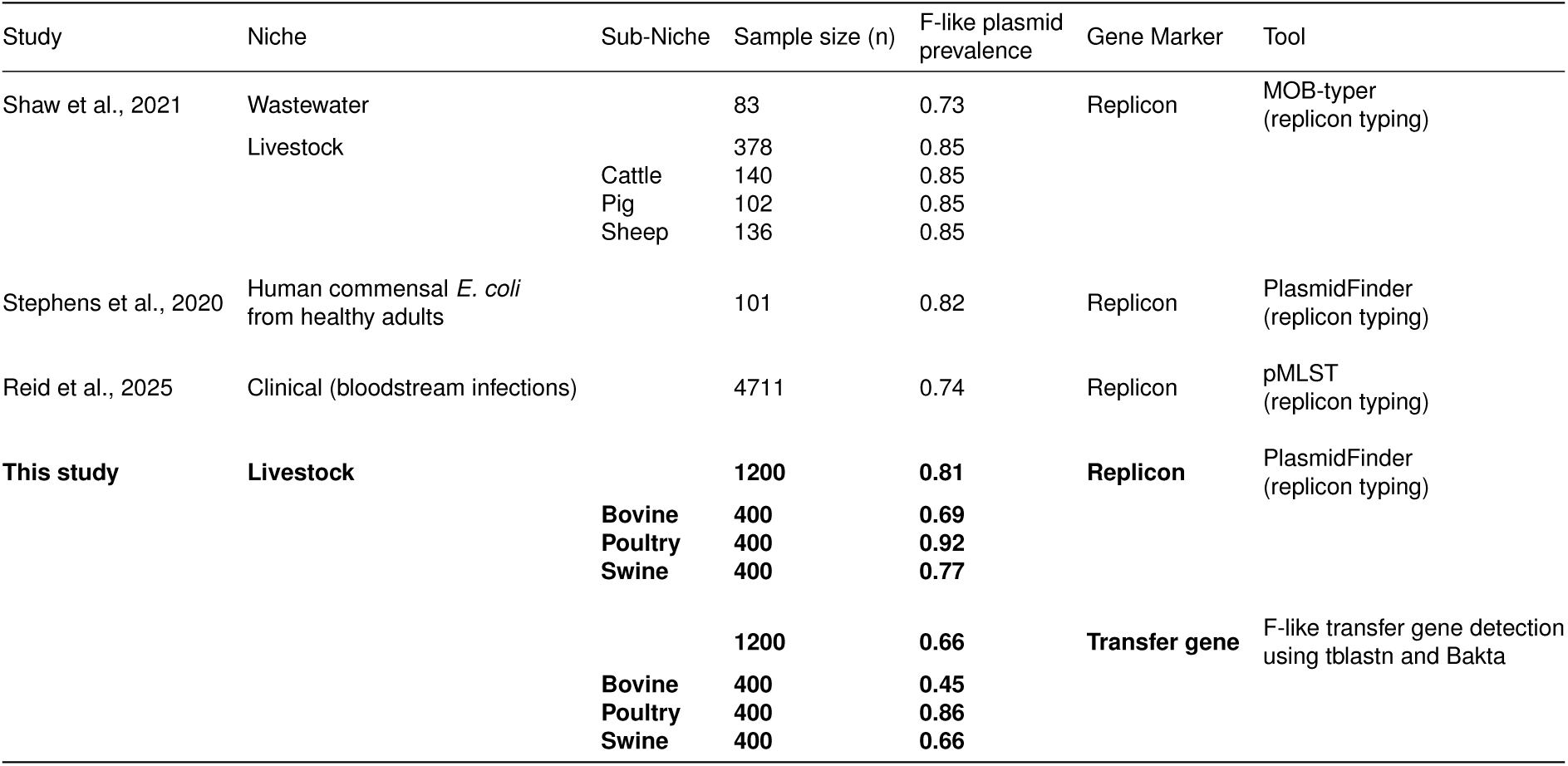
Reported F-like plasmid prevalence estimates for *E. coli* isolated from different environments. In this study, prevalence was estimated using two different types of marker genes: IncF replicons, or transfer genes (here with a detection threshold of at least 5 transfer genes).

### 3.2 Animal environments differ in F-like transfer operon completeness

The inconsistency between the replicon and transfer gene based prevalence estimates suggests that the three animal environments differ in the types of F-like plasmids present, specifically in the completeness of their transfer operons. Completeness of the transfer operon is a proxy for functionality, as plasmids lacking essential transfer genes cannot conjugate on their own [8, 24]. We classified each genome into three completeness categories, based on the presence of 23 F-like transfer gene markers with an identified function in conjugation (Figures 1A, Table S1). Mutations in these genes have been shown to block or reduce transfer [3, 7, 8]. We, henceforth, refer to this set of 23 genes as ”essential genes”. Genomes were classified as ‘missing’ F-like transfer operons if *<* 5 essential genes were detected, ‘incomplete’ for ≥ 5 and *<* 23 essential genes, and ‘complete’ if all essential genes were detected. We note that this classification is not influenced by genome quality: we find no meaningful variation in genome completeness or contamination scores across these groups (Fig S2B,C).

We find a statistically significant association between animal environment and functional completeness (χ^2^(4) = 165*,p <*2×10*^—^*^16^, Cramér’s V effect size = 0.26 [0.22,0.30]). Poultry isolates contained significantly more complete F-like transfer operons (Supplementary Table S2), while bovine isolates were more often missing F-like transfer operons than expected based on independence of variables. Interestingly, the proportion of functionally incomplete F-like transfer operons are similar among the three animal environments.

We compared our classification results to the mobility typing scheme used in Coluzzi *et al.* [22], which relies on the relaxase and a small number of transfer genes to distinguish conjugative, mobilizable and non-mobilizable plasmids (Supplementary Text). In this case, we found the same rank order of prevalence of conjugative plasmids across the three animal environments (Poultry *>* Swine *>* Bovine; Figure S5). However, the absolute prevalences increased because the scheme of Coluzzi *et al.* requires fewer transfer genes to classify a transfer operon as conjugative.

We then assessed whether operon completeness is associated to specific plasmid identities. To do so, we used the MOB-recon tool from the MOB-Suite package to assign plasmid cluster identities to the F-like contigs in each draft genome (hence termed F-like plasmid clusters, Supplementary Text). Only 38.5% of genomes containing an F-like transfer operon (either complete or incomplete, n=786) could be assigned a plasmid cluster. Notably, we find that the same F-like plasmid cluster can be mapped to both functionally complete and incomplete transfer operons (Figure **??**), implying that the same plasmid backbone can differ in their putative ability to conjugate.

To better understand how operon completeness relates to the the total number of transfer genes hits found in each genome, we counted all occurrences of the 35 transfer gene markers in each isolate and compared the distribution of transfer genes across the three environments. Here, multiple occurrences of a transfer gene were counted separately. We find a similar distribution of transfer genes in each animal environment (Figure 2E). The distribution is bimodal, with most isolates either completely lacking an F-like transfer operon or having between 23-35 F-like transfer gene hits. The exception is a peak in Bovine *E. coli* at 3 F-like transfer genes (n = 59 isolates). Interestingly, 34.1% of the isolates with ≥ 23 genes —the size of our essential gene set— have functionally incomplete transfer operons. We also observe a long tail of isolates containing more than 35 transfer genes (9.17% of all isolates), likely indicative of the presence of multiple F-like transfer operons.

We also checked how transfer gene hits were distributed across contigs in a genome and whether there are associations to operon completeness (Figure S4A). Over 66% of genomes with complete F-like transfer operons have transfer genes detected on just one contig. Complete genomes had statistically fewer contigs with transfer genes compared to incomplete genomes (Wilcoxon rank-sum test with continuity correction, W=47564, *p <*2×10*^—^*^16^). We observe strong conservation of gene synteny even when genes are split across multiple contigs, with 92% of genomes (in which gene synteny can be assessed, n=872) containing contigs with transfer genes ordered as found on the F plasmid (S4B).

### 3.3 Differences in plasmid prevalence across animal environments persist after correction for confounders

We next asked whether the difference in prevalence of complete F-like transfer operons across the three animal environments might be confounded by significant differences in *E. coli* phylogenetic background (PhG A-F), sample source (whether the sample came from food or the live animal), presence of antimicrobial resistance genes (ARG presence yes/no), isolate or sampling year (before 2017, 2017 and later) (Figures 3A, S7 and Table S3). We used logistic regression to model the odds of harbouring complete F-like transfer operons with these covariates and their interaction terms (Figure 3B). The model with the lowest AIC included animal environment, PhG, sample source and ARG gene presence as main effects (Table S4) and an interaction between PhG and ARG presence (Table S5). Sampling year is not included as a co-variate, which is additionally supported by the observation that we find no difference in F-like transfer operon completeness in E. coli sampled before 2017 and after 2017 (χ^2^(1) = 3.63*,p* = 0.057, Cramér’s V effect size = 0.05 [0.00, 0.11]), Figure S7)

**Figure 3:**
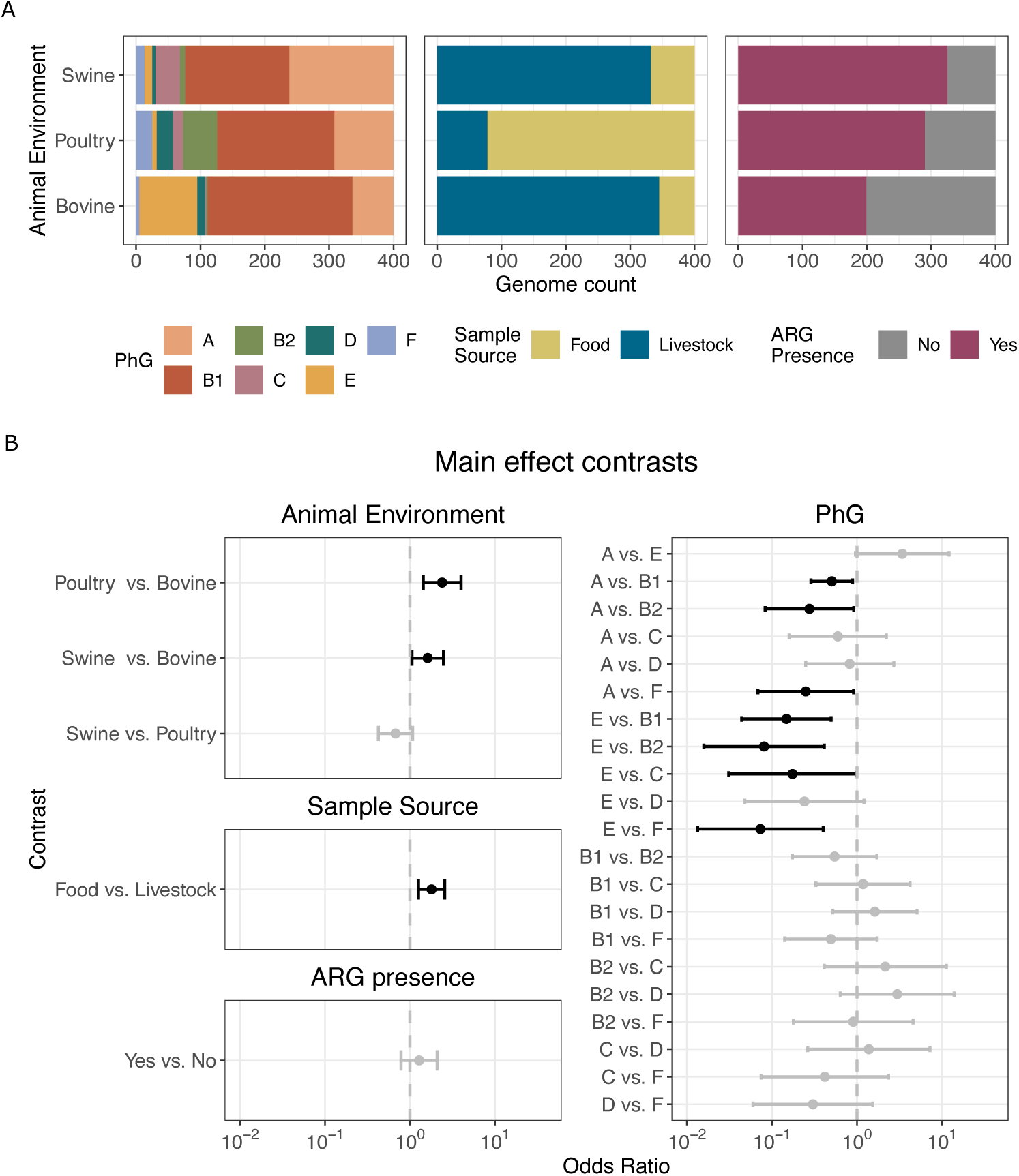
(**A.** Left: *E. coli* phylogroup (PhG) abundances per animal environment. Middle: Proportion of genomes isolated from a given sample source and animal environment. Livestock refers to the live animal, while Food refers to any sample recovered from food products. Right: Presence of at least one antimicrobial resistance gene (ARG) in isolates from a given animal environment. **B.** Odds ratio of harboring a complete F-like plasmid in one group vs. the other after adjusting for all other covariates. The x-axis is on a log scale, and error bars indicate 95% confidence limits. Contrasts that were significant (p *<* 0.05) after correction for multiple testing are depicted in black.

Even after adjusting for these confounders, we find significant differences in the odds of harbouring a complete F-like plasmid across the different animal environments. Both poultry and swine *E. coli* have higher odds than bovine isolates (OR: 2.39 [1.43, 4] and 1.62 [1.06, 2.47] respectively). Swine *E. coli* has lower odds compared to poultry, although the difference is not statistically significant (OR: 0.68 [0.42, 1.08]).

There are also independent effects of the sample source and phylogroup on the odds of harbouring a complete F-like transfer operon. Food isolates have significantly higher odds of harbouring a complete F-like operon compared to livestock isolates (OR = 1.8, 95% CI: 1.26, 2.57), a difference that is not specific to a particular animal environment. Among the different phylogroups, all significant differences involved either PhG A or E (Figure 3D). We find significantly lower odds of a complete F-like operon in PhG A than in B1, B2, or F (0.51 [0.29, 0.89]; 0.28 [0.09, 0.92]; 0.25 [0.07, 0.91], respectively) and significantly lower odds in PhG E than B1, B2, C, or F (0.15 [0.04, 0.5]; 0.08 [0.02, 0.41]; 0.17 [0.03, 0.98]; 0.07 [0.01, 0.4] respectively).

Interestingly, across all genomes we find no significant difference in complete F-like operons between *E. coli* that carry resistance genes and those that do not (Figure 3). Yet this picture changes when we take into account the interaction between resistance carriage and phylogroup (Supplementary Figure S8). Within PhG A, the odds of harbouring a complete F-like operon were significantly higher in isolates carrying resistance genes (2.79 [1.54, 5.04]). In PhG B2 instead, the odds were significantly lower in isolates carrying resistance genes (0.17 [0.04, 0.69]).

A sensitivity analysis repeating the logistic regression procedure on a dereplicated dataset (1,160 genomes; excluding isolates with ≥ 99.99% ANI, identical sampling metadata, and similar plasmid content; Figure S3C,) confirmed that most observed effects remained robust. However, we note that the difference between swine and bovine isolates falls below statistical significance (Figure S9A).

Therefore, we show that animal environment differences in F-like operon presence remain significant even after accounting for confounding by phylogroup, presence of resistance and sample source.

### 3.4 Transfer gene frequencies are correlated with gene function

We asked whether differences in F-like transfer operon prevalence between animal environments extend to more fine-grained differences in transfer gene presence/absence

between animal environments. We estimated individual gene frequencies for each of the 35 transfer genes relative to the total number of genomes with at least one transfer gene in a given environment (Figure 4). This is to avoid inflating gene frequency differences due to the different baseline frequency of F-plasmid presence in different environments. Even with this definition, the median gene frequencies differ by animal environment (62% in bovine, 81% in swine, and 91% in poultry respectively), reflecting differences in functional completeness across the environments. To focus on the prevalence of specific genes relative to the rest of the operon, we interpret patterns of transfer gene content relative to the median gene frequency of the transfer operon in that animal environment. Frequencies at or higher than the environment-specific median gene frequencies suggest that the gene is important in the conjugation process, whereas frequencies lower than the environment-specific median suggest non-essentiality.

**Figure 4:**
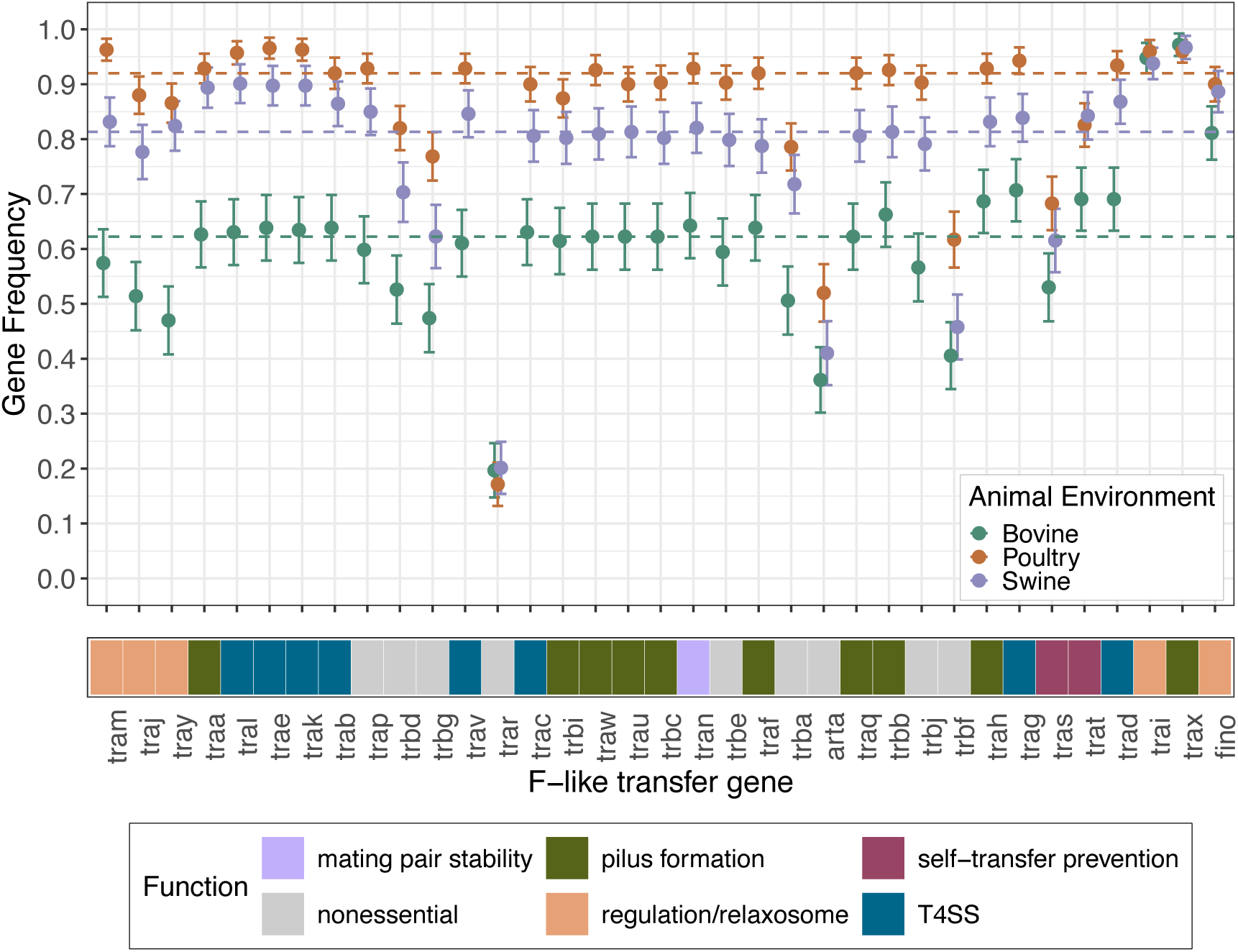
Gene frequencies of individual transfer genes in *E. coli* genomes isolated from bovine (green), poultry (orange), and swine (purple). Transfer genes are coloured by their role in conjugation. Gene frequencies were calculated relative to the total number of genomes from each animal environment (n=400) that contained at least one transfer gene hit. The bars indicate 95% confidence intervals, computed using a normal approximation to the binomial distribution. The coloured dashed lines refer to median gene prevalences within each animal environment.

We find that the relative transfer gene frequencies are mostly independent of the animal environment, and are primarily determined by their functional role in conjugation. Genes previously described as essential for conjugation have higher frequencies than those that are non-essential (Figure 4). The relaxase *traI* is present at frequencies over 90% in all three animal environments, indicative of its important role in conjugation and mobilization. The other pilus formation genes and genes involved in the type IV secretion system (T4SS) have gene frequencies close to or higher than the median gene frequency in each animal environment (Figure S10). Instead, non-essential genes typically have frequencies lower than the median gene frequencies, indicating that these genes are often missing in operons. Two exceptions are *traP* and *trbE*, non-essential genes with frequencies closer to the median gene frequencies in each niche, hinting that they may play a functional role in natural environments. Transfer gene *traR*, described to be non-essential, has the lowest gene frequencies in all three animal environments, at around 19%.

Interestingly, regulatory genes like *traM*, *traJ*, *traY*, and *finO* and self-transfer prevention genes *traS* and *traT* have variable patterns of presence in the three animal environments (Figure 4, Supplementary Figure S10), potentially indicating environment-specific diversity patterns.

### 3.5 Poultry *E. coli* host a larger diversity of conjugative plasmids

Since *E. coli* often carry multiple plasmid types simultaneously [54], we asked whether we also find differences in the prevalence of non-F conjugative systems between the animal environments. To do so, we identified which mating pair formation (MPF) types are present in each isolate using geNomad [48]. Because MPF types are defined based on sequence homology of components of the T4SS and other proteins involved in transfer, they directly reflect the types of conjugative plasmids present in the isolate. We find that more than a third of the isolates (35.91%) carry more than one MPF type (Figure S11).

We found significant differences between the animal environments in the number of non-F MPF types carried per isolate (χ^2^(2) = 66.22, *p <* 0.001, Kruskal–Wallis rank-sum test). In terms of carriage, poultry *E. coli* harbour significantly more non-F MPF types than bovine and swine *E. coli* (Figures S12, 5A). Swine and bovine do not differ significantly in their number of non-F MPF types. Interestingly, in both poultry and swine, we find no difference in non-F MPF type carriage between isolates that carry complete, incomplete or lack F-like operons (χ^2^(2) = 2.72, *p* = 0.26, χ^2^(2) = 1.02, *p* = 0.60 respectively in poultry and swine). Interestingly, bovine isolates missing F-like operons seem to carry fewer plasmids of other types too (missing vs. incomplete *p <* 0.01; missing vs. complete *p <* 0.05). Generally, however, the presence of F-like plasmids does not seem to influence the number of other conjugative plasmids carried by these *E. coli* isolates.

We looked at the prevalences of the individual MPF types across the animal environments (Figure 5B). In addition to the significant differences in prevalences of MPF F across animal environments, we also find significant differences in the prevalence of MPF I (χ^2^(2) = 75.94, *p <* 0.001) and MPF G (χ^2^(2) = 11.323, *p <* 0.01). Poultry isolates have a higher prevalence of MPF I than swine and bovine isolates. This higher prevalence of MPF I was the driver of overall higher numbers of non-F MPF types in poultry (Figures 5A, S12).

**Figure 5:**
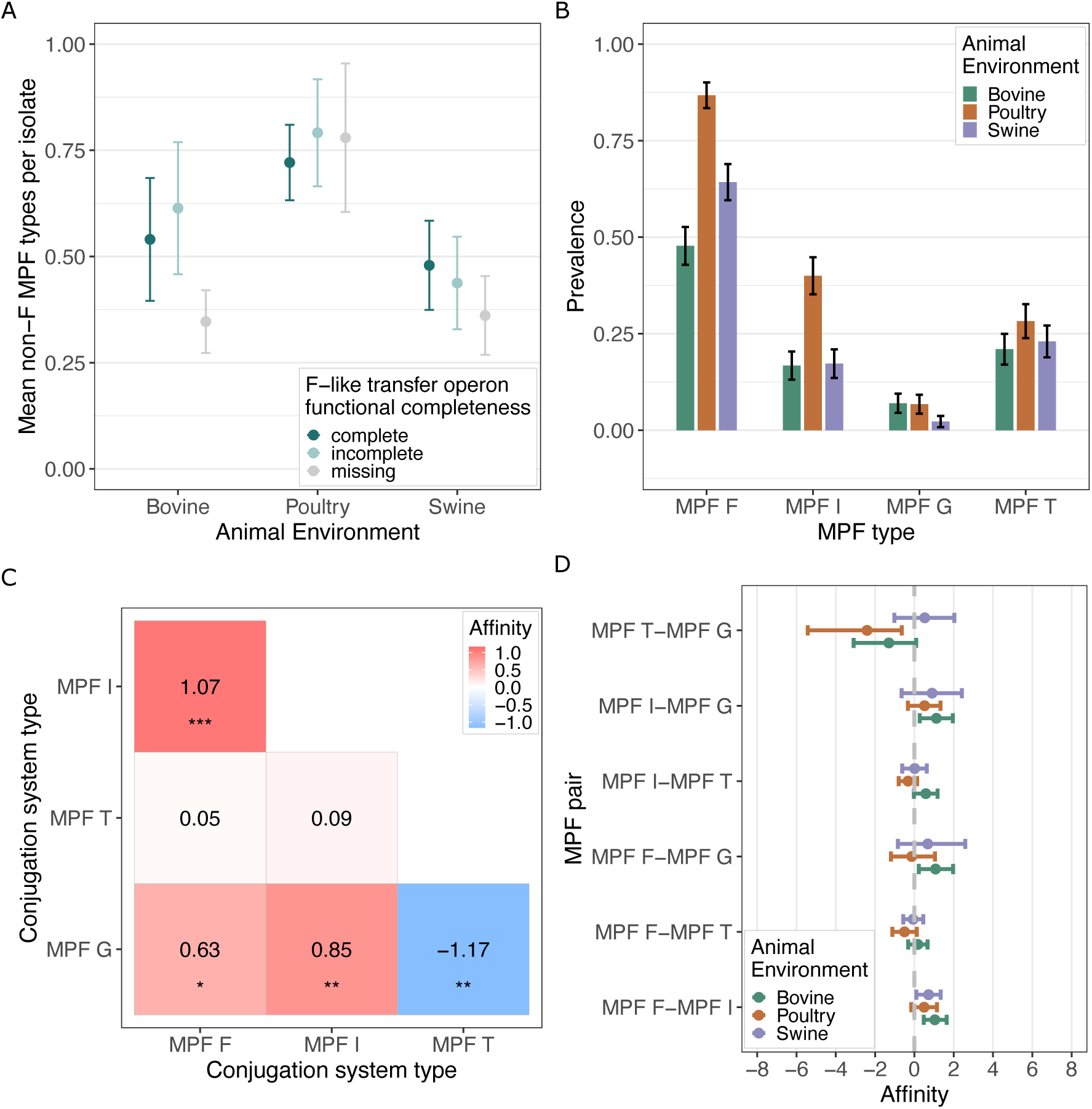
**A.** Mean number of non-F MPF types per isolate across animal environments and F-like completeness categories. Error bars are 95% confidence intervals around the mean. **B.** Prevalences of MPF types across animal environments. MPF prevalence is computed as the proportion of genomes where at least 3 distinct genes specific to an MPF type were detected. The error bars indicate 95% confidence intervals, computed using a normal approximation to the binomial distribution. **C.** Affinity matrix describing the odds of finding pairs of conjugation system types together in a isolate. Positive values (red) indicate higher odds of finding a pair together than the null expectation of random assortment. Negative values (blue) indicate higher odds of finding a pair in two different isolates than the null expectation. A 0 value (white) indicates co-occurrence as expected based on individual frequencies (null expectation). Significance levels: *p ≤ 0.05, **p ≤ 0.01, ***p ≤ 0.001. **D.** Affinity of MPF pairs across animal environments.

### 3.6 Co-occurrence patterns of conjugative systems are similar in all three environments

We wanted to test whether specific conjugative plasmid types co-occur with F-like conjugative plasmids in an isolate and whether these patterns differ by animal environment. We assessed whether pairs of MPF types are detected together in an isolate more or less frequently than expected based on the prevalence of individual MPF types and a null distribution of independent random assortment [55](Supplememtary Text). As a test of our method, we first assess the co-occurrence between MPF and relaxase types in the entire dataset. Since a functional conjugation system requires both an MPF system and a relaxase, we expect to find mostly positive associations between MPF and MOB types. Indeed, we find 12/20 MPF-MOB pairs that are significantly positively associated (Figure S13).

We next assessed co-occurrence between pairs of MPF types in the entire dataset (Figure 5C), focusing here on the associations with MPF F. Plasmid types MPF I and MPF G co-occur more often with MPF F than expected by chance. On the other hand, MPF T is not positively or negatively associated with MPF F. These patterns also persist at the level of the individual animal environments (Figure 5D). This suggests that regardless of the environment, similar conjugative plasmids tend to be found together with F-like plasmids. In fact, patterns of co-occurrence are similar across the animal environments for almost all pairs of MPF types assessed.

## 4 Discussion

In this work, we investigated the genetic diversity of F-like transfer operons in *E. coli* from livestock environments, to understand the distribution of transfer potential in natural bacterial populations. We used F-like transfer gene presence to measure transfer operon completeness as a proxy for transfer potential. This novel approach showed that livestock environments differ in their prevalence of complete F-like transfer operons, with a significantly higher frequency found in poultry (56%), than swine (36%) or bovine (22%) environments (Figure 2D). Notably, this difference in transfer potential persists even after accounting for possible confounders such as *E. coli* phylogenetic background, presence of antibiotic resistance genes and isolation from food or the live animal (Figure 3B, S8).

While many studies use the replicon to estimate plasmid presence in natural environments (e.g. [30–33]), we find that replicon-based plasmid prevalences are systematically higher than our transfer-operon based prevalence estimates. This suggests that studies that rely only on replicons to assess plasmid prevalence may overestimate the transfer potential of F-like plasmids in natural environments. Indeed in our study, 19.6% of the isolates containing an F-like replicon were missing F-like transfer operons entirely and 32.42% were lacking essential genes required for self-conjugation (Figure 2C).

Our estimates of functional F-like plasmid prevalence provide insight into the distribution of plasmid mobility at the population level and how this varies by environment. Transitions in plasmid mobility are thought to be frequent, mostly involving deletions or pseudogenizations of transfer genes, leading to a loss of the ability to self-conjugate [22, 23]. While we observe differences in the frequencies of genomes with complete or entirely missing F-like transfer operons, the frequency of incomplete F-like operons are similar across the three environments (∽25%; Figure 2D). This suggests that the per-plasmid rate of transitioning from complete to incomplete or incomplete to missing differs across the three host environments. To understand how different environments can maintain different frequency distributions of the three mobility classes, it will be important to model selection on mobility and the rate of transitions between classes.

This study also sheds light on the relative roles of environment and phylogenetic background in shaping plasmid diversity. In our collection of livestock-associated *E. coli*, we found that the variance in F-like transfer potential is better explained by the animal environment than the *E. coli* phylogenetic background (Figure 3B). This is consistent with *E. coli* sampled from diverse environments in Australia [51], where mobile genetic element-associated gene variation was explained more by environment than *E. coli* phylogroup. However, it contrasts with what has been observed at broader taxonomic scales. In a study on plasmids from diverse bacterial isolates across multiple environments, bacterial host phylum explained more variation in plasmid relaxase genes than the isolation environment [35]. The distribution of plasmid mobility has also been found to vary across bacterial taxa, independent of the environment they are found in [22]. These observations suggest that at broader phylogenetic scales, lineage-specific constraints most strongly dictate plasmid diversity, whereas at the level of a single bacterial species environment-related effects can become more apparent. Further comparative studies within bacterial species other than *E. coli* will be necessary to determine how general this scale-dependent pattern is.

We find that poultry isolates are distinct from both swine and bovine isolates, hosting more conjugative plasmids per isolate (Figures 5A,B, S12). Our observation is consistent with studies that reported enrichment of MGEs in poultry isolates [51, 56]. In addition to this enrichment in MGEs, poultry isolates have also been found to be enriched in virulence and resistance genes [51]. This stand-alone position of poultry could be due to the specific selection-pressures found in poultry farming, where heavy antibiotic use is often the norm [56], or inherent differences in animal physiology [57] and microbiome [58, 59]. Distinguishing between these different hypotheses would require targeted sampling of poultry farms coupled with farm-specific metadata on antibiotic consumption.

Despite differences in the frequencies of conjugative plasmids between the environments, the patterns of transfer gene presence and absence are similar (Figures 4,S10) and correlates with functional importance in conjugation. This suggests that selection for a given transfer gene is similar across the environments. We find exceptions to this pattern in essential transfer genes involved in regulation and exclusion, which display the considerable variability in gene frequencies across environments, suggesting different environments might have different preferred regulatory variants and exclusion potential. A previous study looking at F-like plasmid transfer operons in *Enterobacteriaceae* across a diversity of environments found alternate regulatory configurations in non-*E. coli* species [24]. That we find such patterns also in a single bacterial species suggests that the regulation of conjugation is also modulated by environment-specific selection pressures, and not solely constrained by phylogenetic background.

Our gene-based detection protocol assumes that the presence of F-like transfer genes in the genome is a sufficient condition for the formation of a conjugation apparatus, even if the genes forming the operon are on different contigs. This assumption is supported by genetic complementation studies, which demonstrate a plasmid’s nonfunctional transfer gene can be rescued by introducing a functional variant on a second plasmid [60–62]. Thus, a functional conjugation system can still assemble as long as the necessary genes are present and functional, even if the genetic components are on different plasmids. A recent study has also shown that co-resident pED208 and F plasmids (both F-like plasmids) missing essential transfer genes can exchange subunits to build a functional chimeric conjugation system [63]. Our approach demonstrates the viability of draft genomes — which make up the vast majority of public genome databases — as a resource for environmental plasmid surveillance.

Together, our results demonstrate that the environment is a major driver of variation in F-like plasmid prevalence and mobility potential across *E. coli* populations. While our analyses focused on F-like plasmids, they are easily transferable to other mobile genetic elements. Such careful studies of plasmid mobility at the level of individual transfer genes will be essential to arrive at environment-specific assessments of transfer potential, allowing us to predict how plasmids spread in natural bacterial populations.

## Acknowledgments

The authors thank João Pires, Aswin Krishna, Ricardo León-Sampedro and Aravind Srinivasan for helpful comments at various stages of the research and writing process.

## 6 Author contributions

Sneha Sundar: Conceptualization, Methodology, Formal analysis, Writing - Original Draft, Software, Validation, Investigation, Data Curation, Writing Review & Editing, and Visualization

Sebastian Bonhoeffer: Conceptualization, Methodology, Writing - Review & Editing, Supervision, Funding acquisition

Jana S. Huisman: Conceptualization, Methodology, Writing - Original Draft, Supervision, Data Curation, Writing Review & Editing, Visualization, and Funding Acquisition

## S1 Supplementary Figures

**Figure S1:**
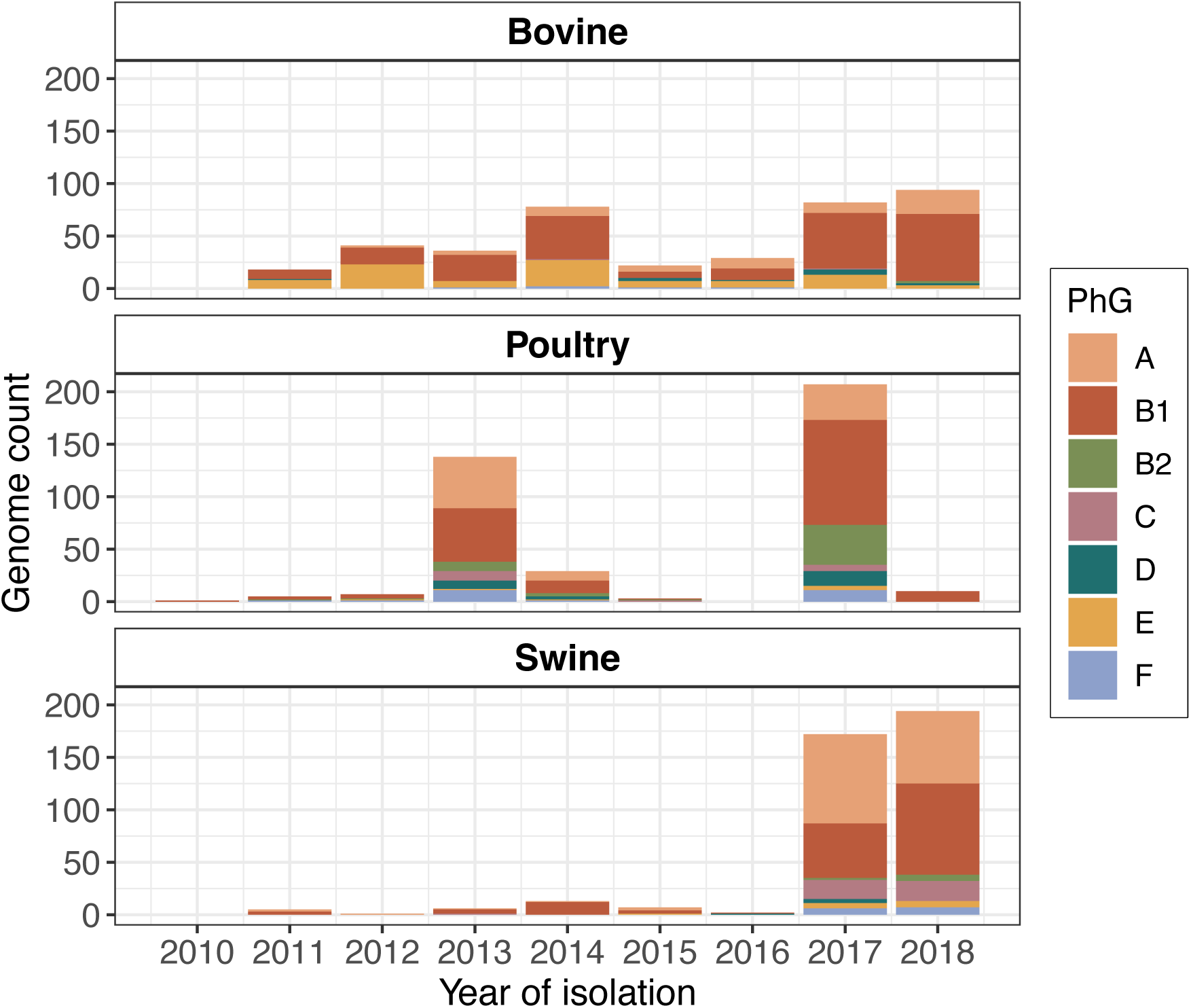
The distribution of our 1200 *E. coli* genomes across animal environments (row; n=400 per animal), isolation date (x-axis), and phylogroup (color).

**Figure S2:**
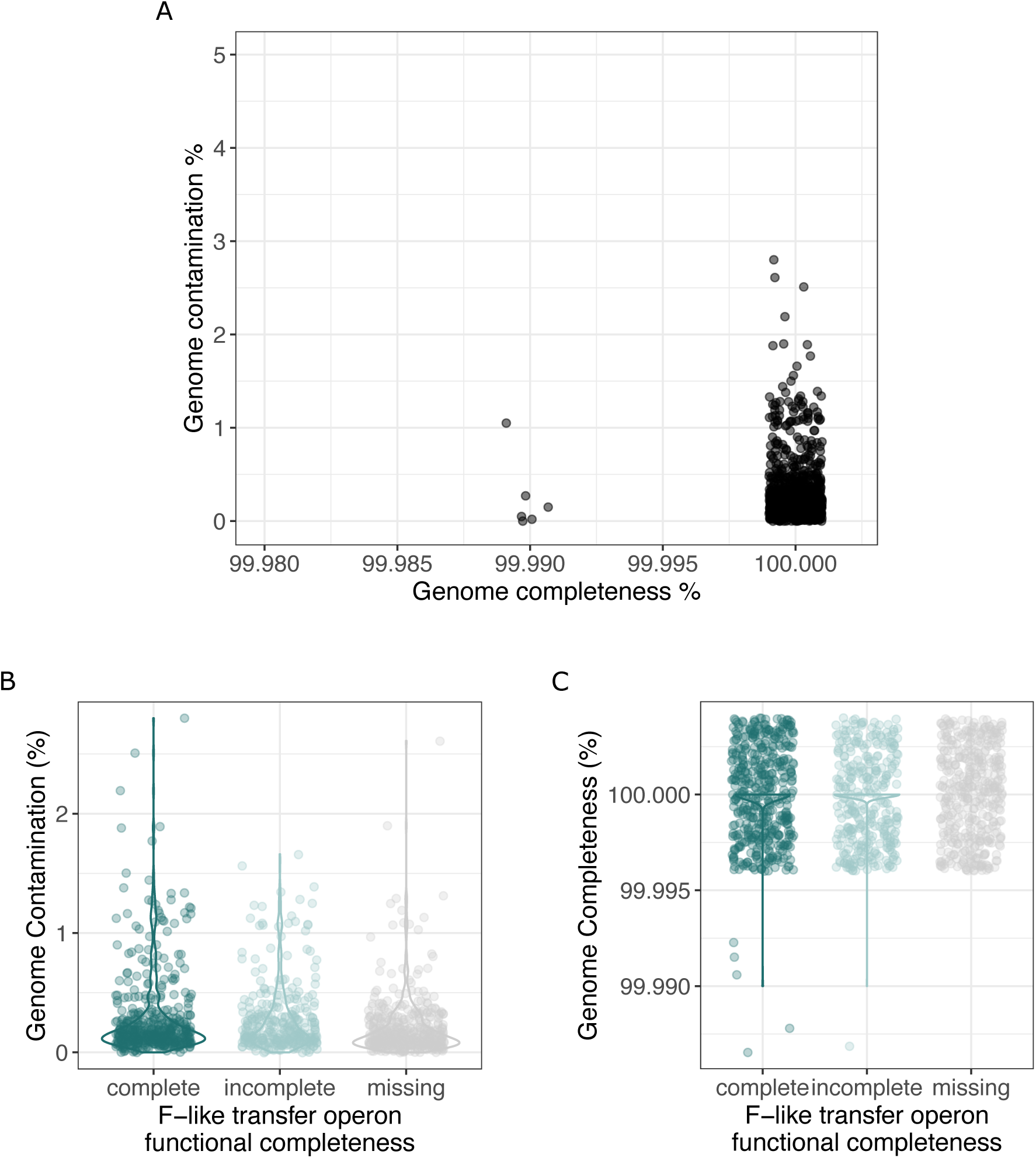
**A.** Genome completeness and contamination scores of all genomes assessed using CheckM2 [1]. All genomes had ≥99.9% completeness and *<*3% contamination and meet the standard criteria for high quality draft genomes (≥95% completeness, ≤5% contamination [2]) **B.** Contamination scores for genomes with complete, incomplete and missing F-like transfer operons **C.** Completeness scores for genomes with complete, incomplete and missing F-like transfer operons

**Figure S3:**
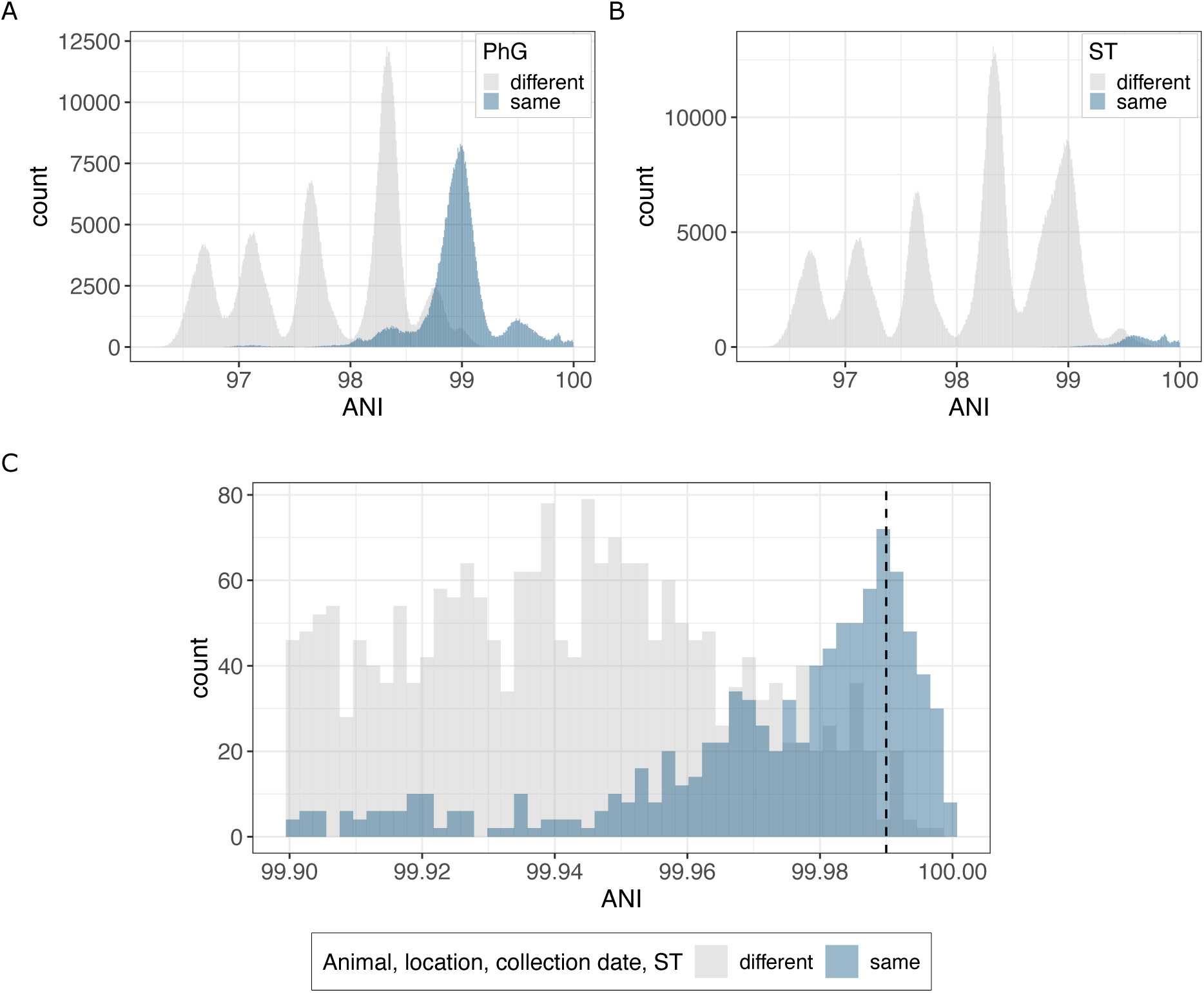
Pairwise Average Nucleotide Identity (ANI) distributions. **A.** Intra-phylogroup (blue) vs. inter-phylogroup (grey) pairs. **B.** Intra-sequence type (blue) vs. inter-sequence type (grey) pairs. **C.** Highly similar pairs (ANI ≥ 99.9%) sharing identical animal environments, locations, collection dates, and sequence types. The dashed line represents the recommended threshold (99.99%) for strain-level classification [3, 4].

**Figure S4:**
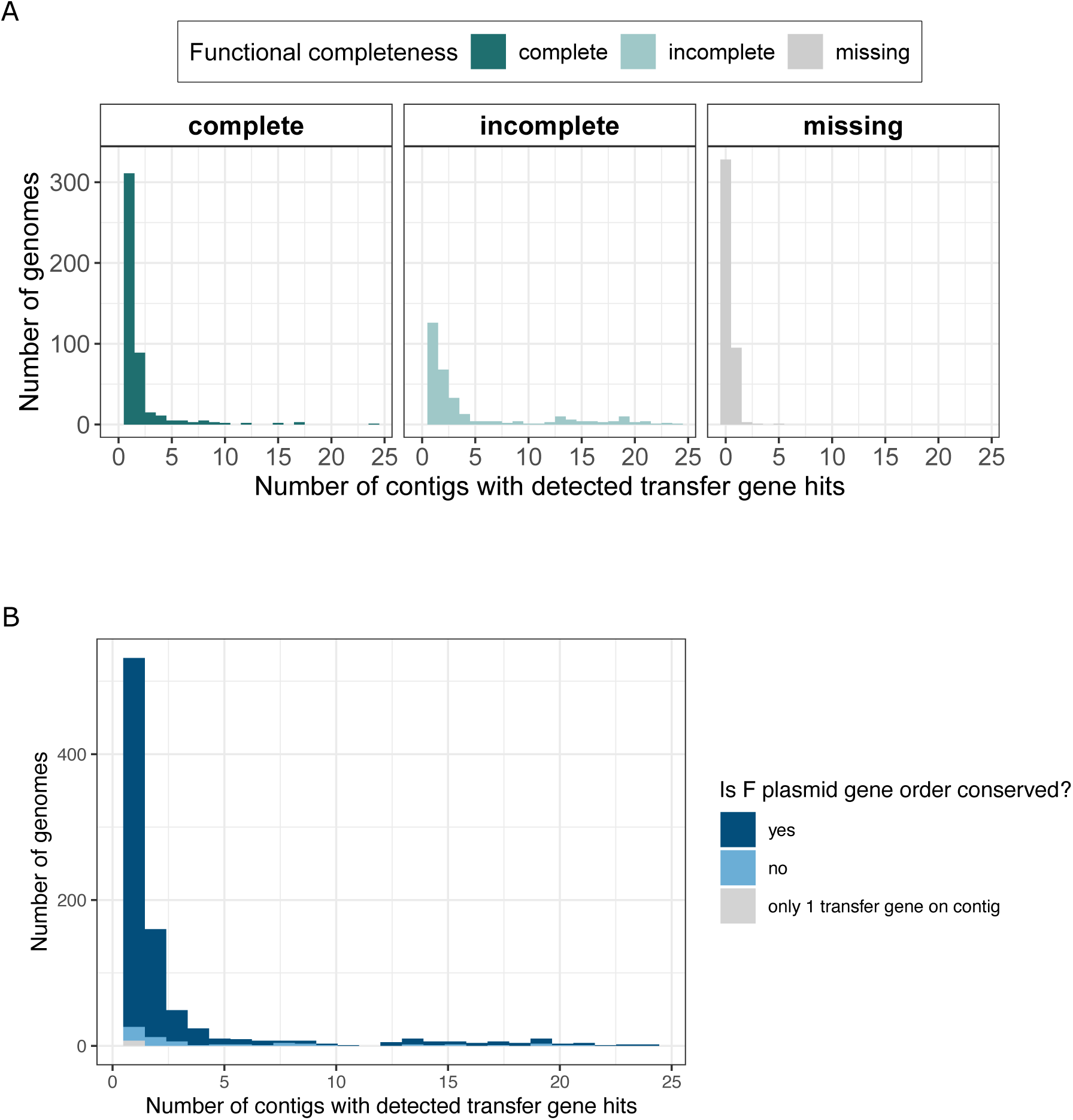
**A.** Distribution of the number of contigs in which transfer genes are detected in genomes with at least 1 detected transfer gene (n = 872). Colour indicates the proportion of genomes in which F plasmid gene order is conserved in all contigs with at least 2 transfer gene hits. **B.** Distribution of the number of contigs with detected transfer genes within each functional completeness category (all 1200 genomes assessed).

**Figure S5:**
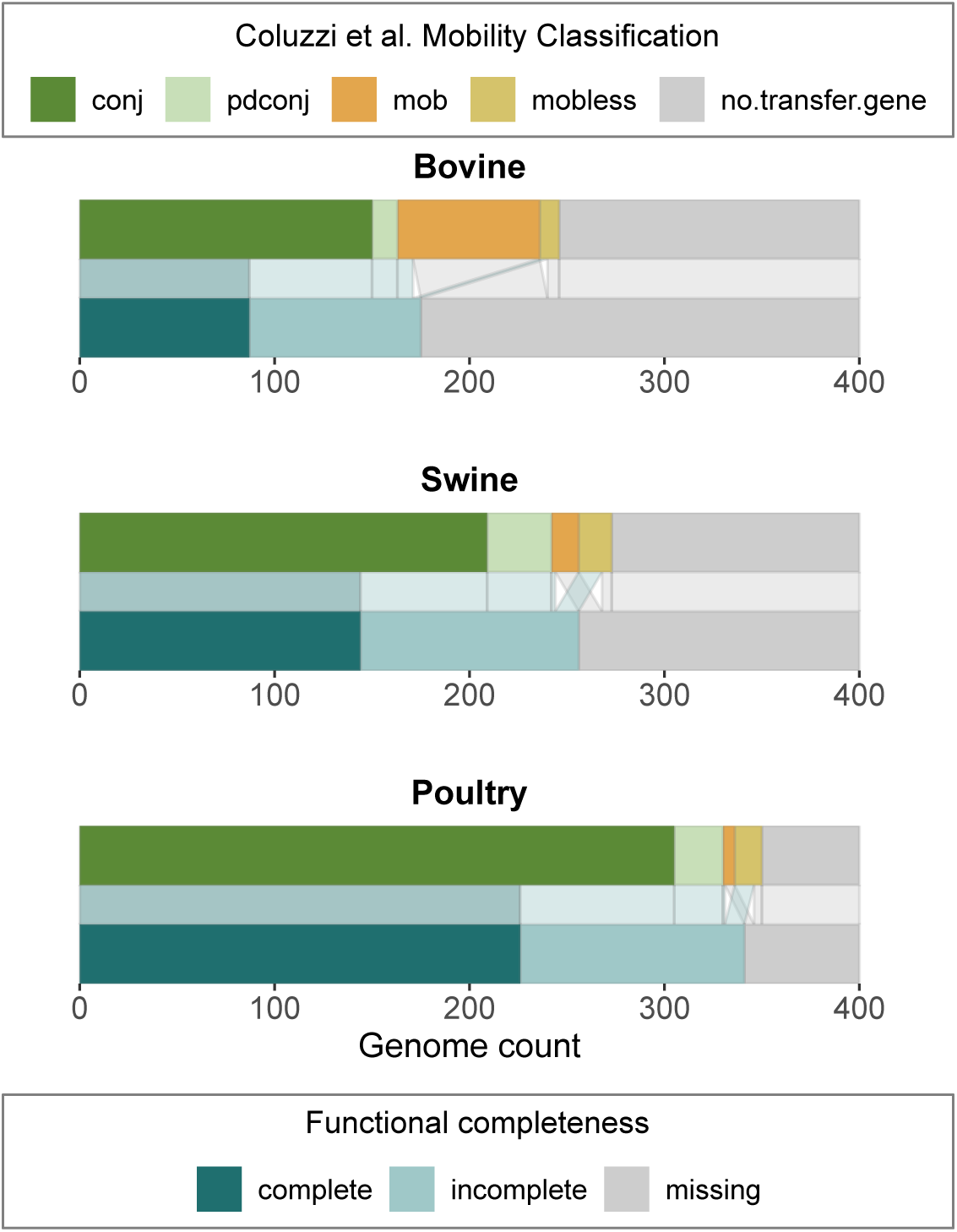
Proportion of *E. coli* isolates with F-like transfer operons within each animal environment, as estimated using either the mobility typing scheme of Coluzzi et al. [5] (top row in each panel) or the functional completeness metric defined in this study (bottom row). Mobility typing in Coluzzi et al. uses the presence of the relaxase and a minimum number of other conjugation-related genes to classify a plasmid: conjugative (conj: contains relaxase, VirB4, T4CP and at least 3 MPF proteins), mobilizable (mob: contains relaxase but missing the required number of other components to be conjugative), mobless (lacking the relaxase) or partially decayed conjugative (pdCONJ: a subcategory of MOB plasmids with more than 5 conjugation proteins excluding the relaxase) or as having no F-like transfer gene (no.transfer.gene). The functional completeness categories used in this study were based on a set of 23 essential transfer genes: ‘complete’ (all 23 essential F-like transfer genes), ‘incomplete’ (between 5 and 22 essential genes) or ‘missing’ (less than 5 essential genes).

**Figure S6:**
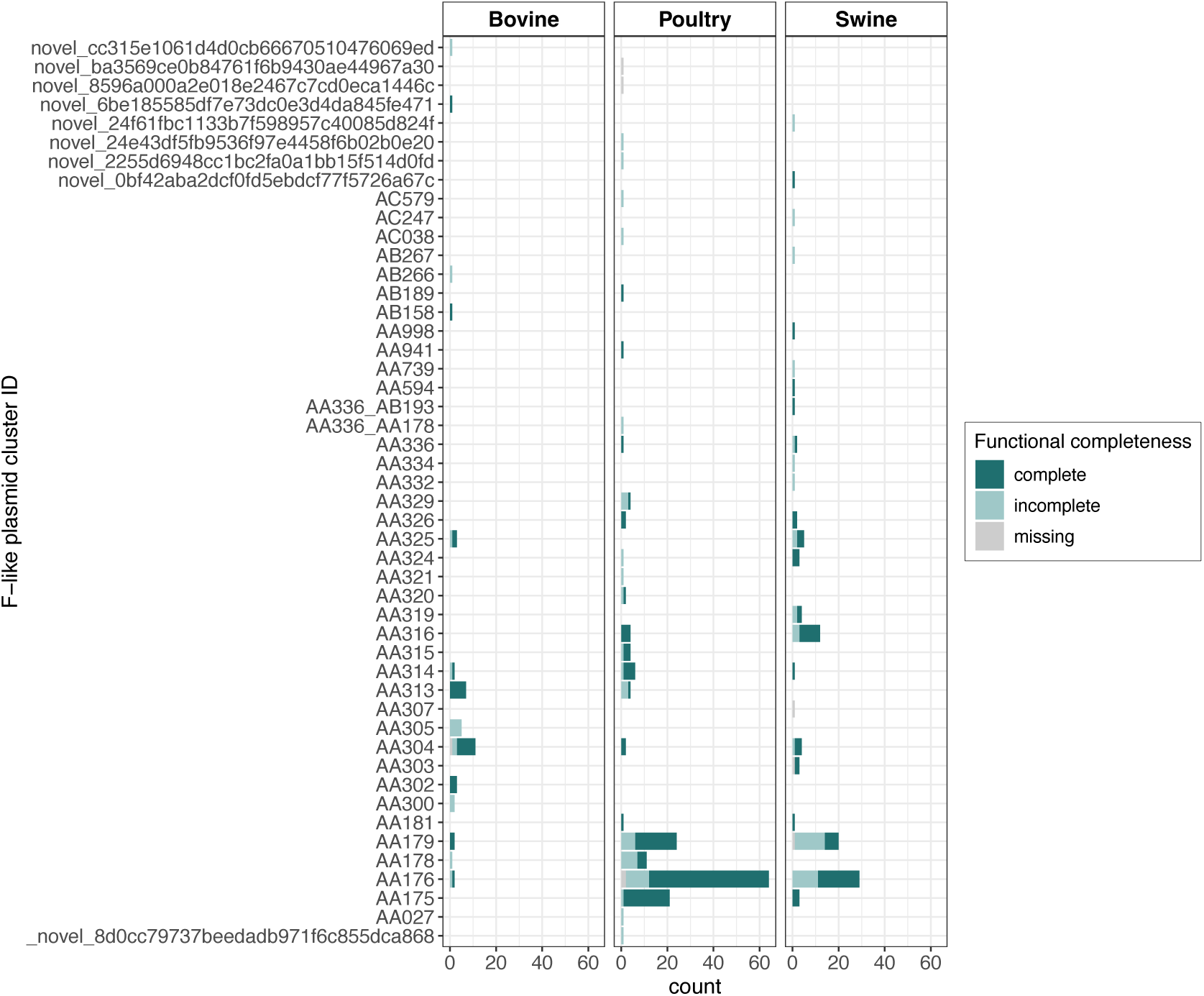
Distribution of F-like plasmid clusters in each animal environments. Bars indicate the proportion of each plasmid cluster classified as having a functionally complete F-like transfer operon (dark blue), incomplete F-like transfer operon (light blue) or missing an F-like transfer operon (grey). Plasmid clusters were assigned using MO-Brecon from the MOBSuite package (Supplementary Text).

**Figure S7:**
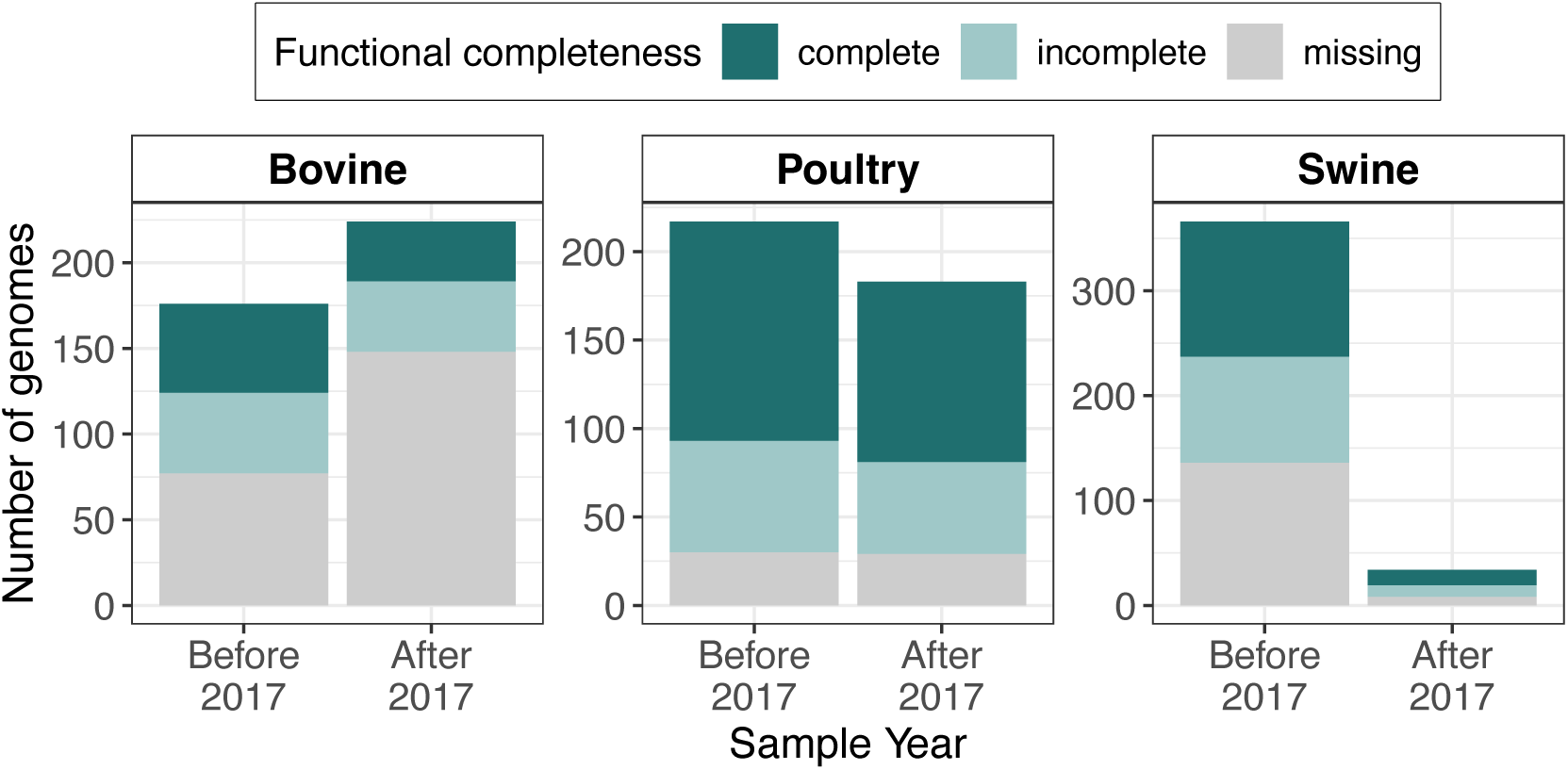
Proportion of *E. coli* isolates in each animal environment with complete, incomplete and missing F-like transfer operons sampled before 2017 and after 2017 (which also includes 2017). We find no difference in F-like transfer operon functional completeness in isolates sampled before and after 2017 (χ^2^(1) = 3.63*,p* = 0.057, Cramér’s V effect size = 0.05 [0.00, 0.11])

**Figure S8:**
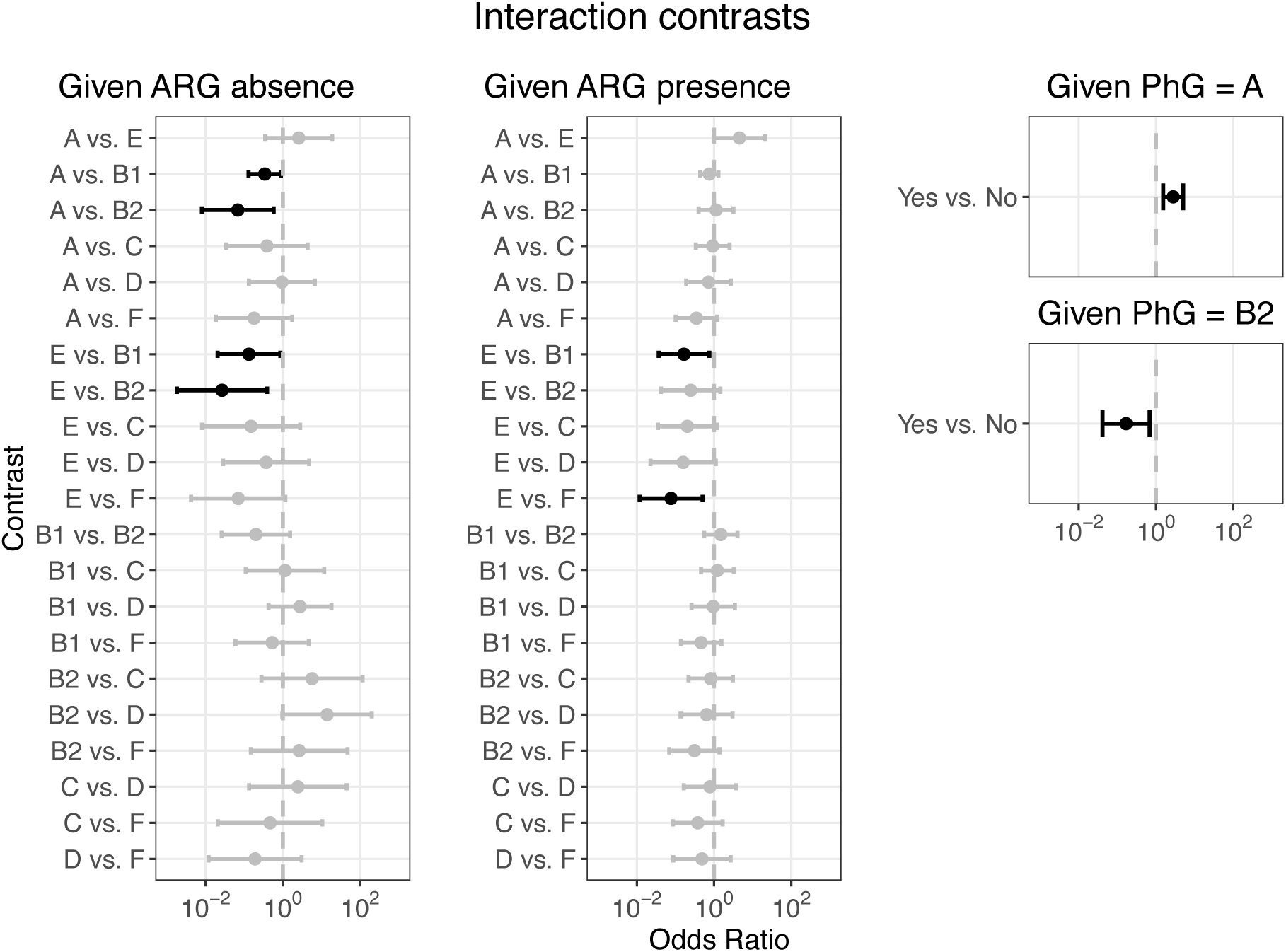
Contrasts for the interaction term between antibiotic resistance gene (ARG) presence and *E. coli* phylogroup (PhG). We report odds ratio of harboring a complete F-like plasmid in one group vs the other conditioned on a specific variable. The x-axis is on a log scale, and error bars indicate 95% confidence limits. Contrasts that were significant (p *<* 0.05) after correction for multiple testing are shown in black, while nonsignificant contrasts are grey.

**Figure S9:**
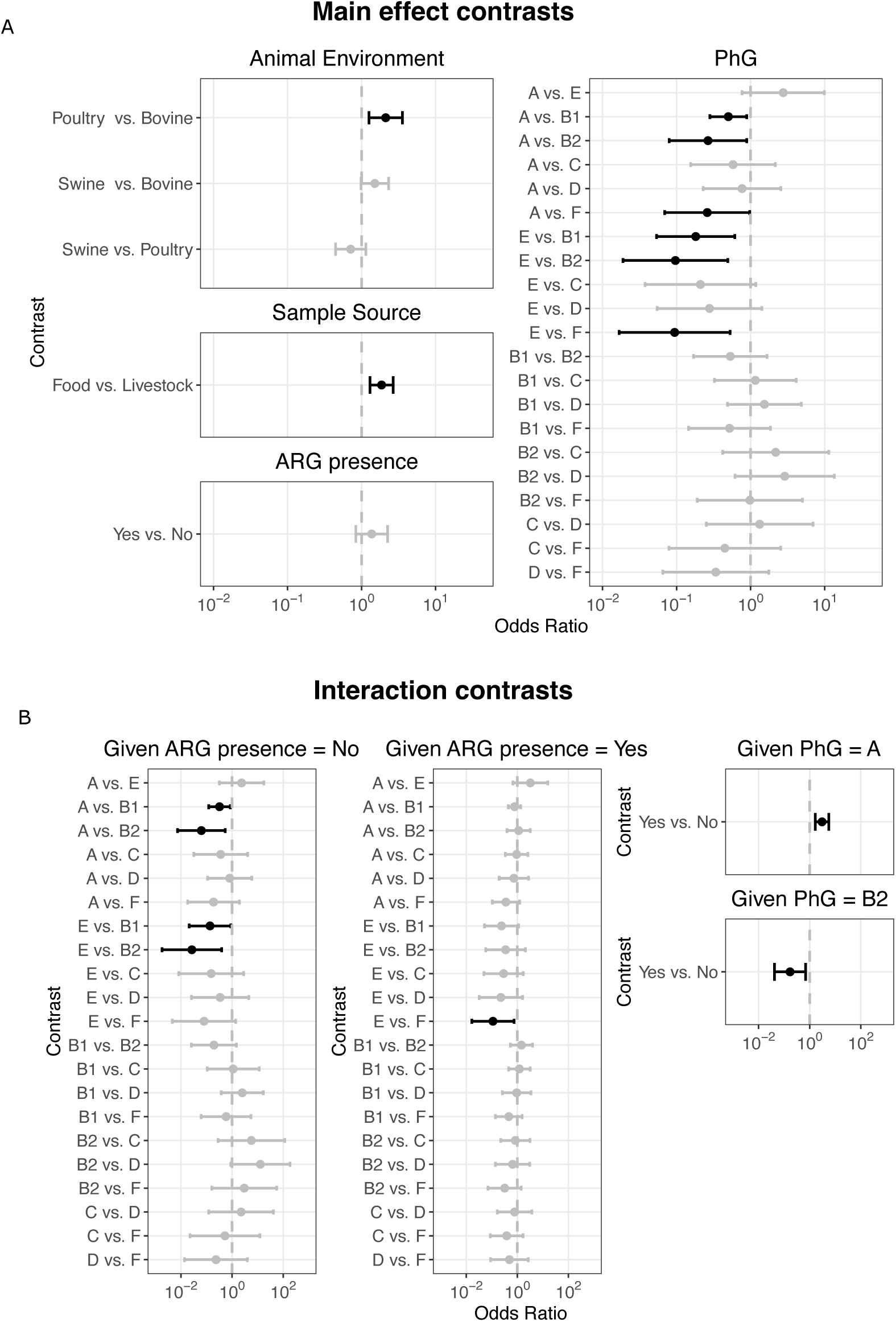
Logistic regression analyses using a reduced dataset of 1160 genomes. The reduced dataset was obtained after removing highly similar genomes (≥99.99% ANI) with similar plasmid content, which were sampled at the same location and time. The model selection procedure was repeated for this dataset and the same covariates were selected. **A.** Main effect contrasts of harboring a complete F-like plasmid in one group vs. the other after adjusting for all other covariates. **B.** Contrasts for the interaction term between antibiotic resistance gene (ARG) presence and *E. coli* phylogroup (PhG). Odds ratios of harboring a complete F-like plasmid in one group vs the other conditioned on a specific variable are reported. For both A and B, The x-axis is on a log scale, and error bars indicate 95% confidence limits. Contrasts that were significant (p *<* 0.05) after correction for multiple testing are depicted in black.

**Figure S10:**
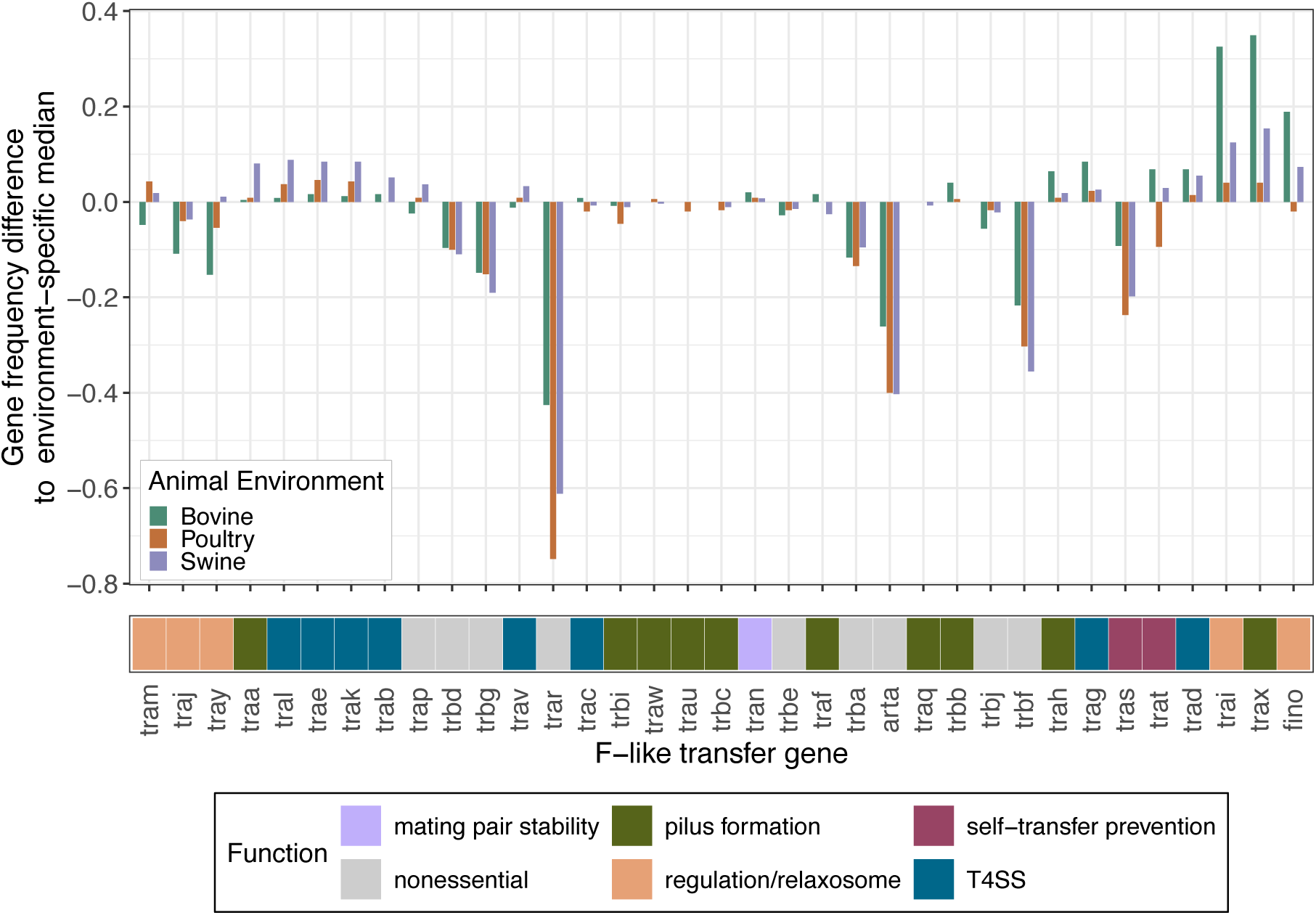
The difference between the frequency of each F-like transfer gene and the median gene frequency of the entire operon within each animal environment: bovine (green), poultry (orange), and swine (purple). Transfer genes are colored by their role in conjugation.

**Figure S11:**
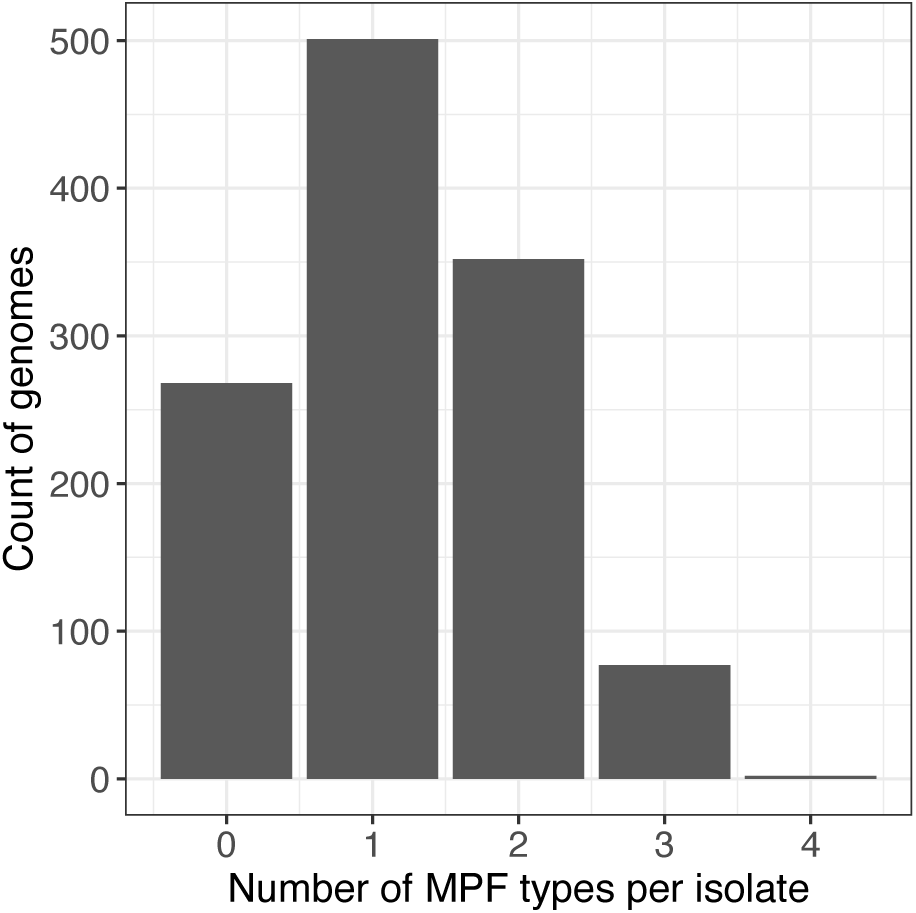
Distribution of the number of MPF types detected per isolate using geNo-mad.

**Figure S12:**
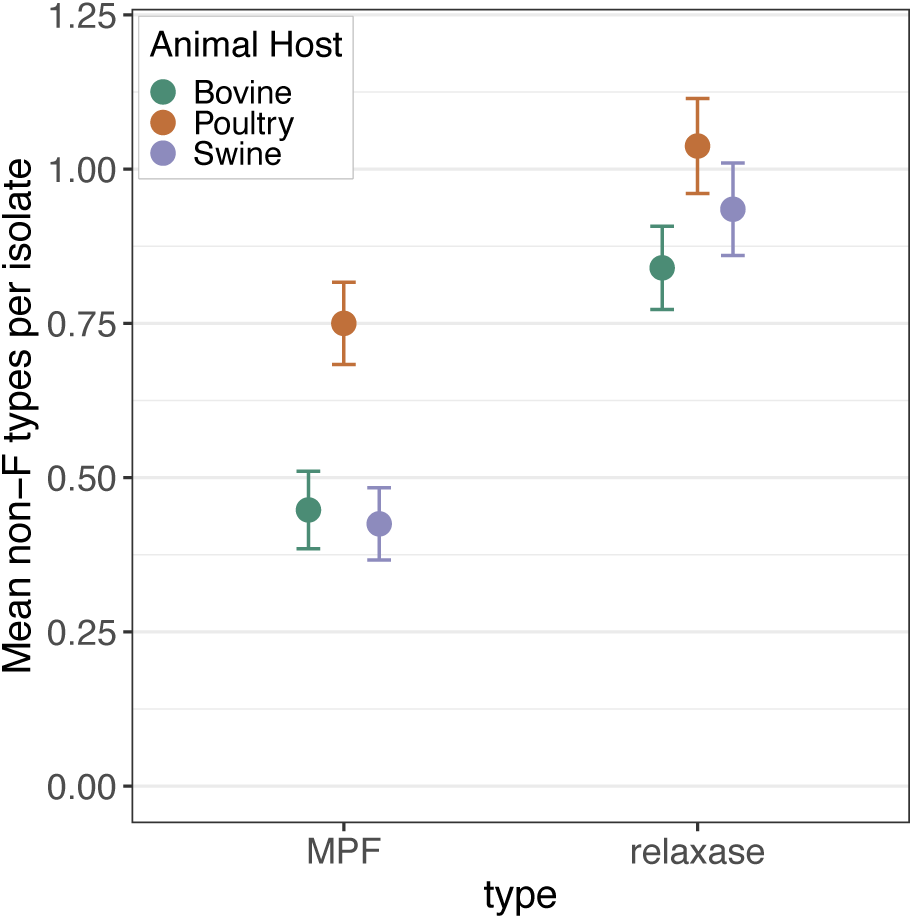
Mean number of non-F MPF types and relaxase types per isolate in each animal environment. MPF and relaxase types were detected using geNomad. Error bars indicate 95% confidence intervals of the mean.

**Figure S13:**
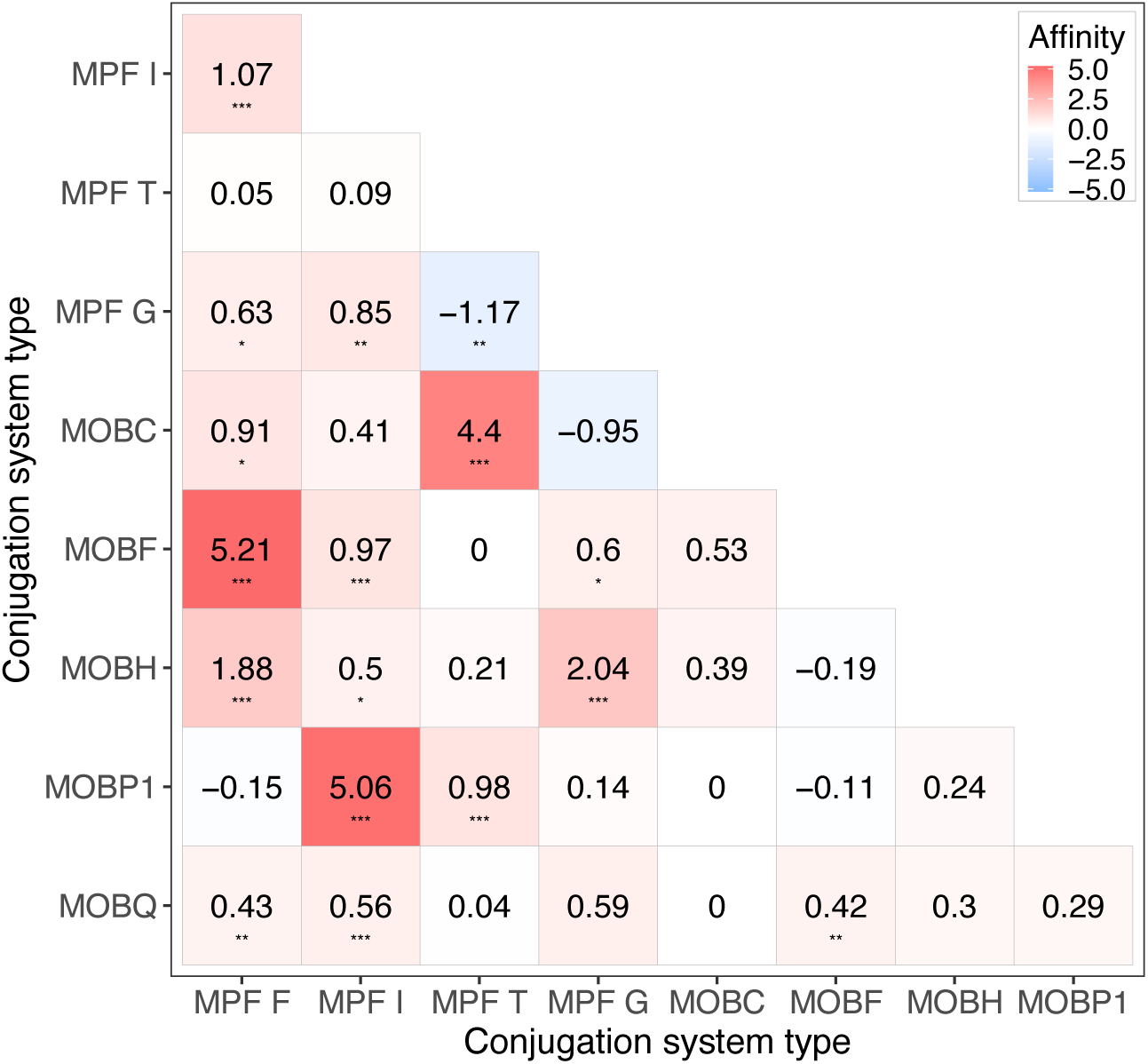
Affinity matrix describing the odds of finding pairs of conjugation system and relaxase types together in a isolate. Positive values (red) indicate higher odds of finding a pair together than the null expectation of random assortment. Negative values (blue) indicate higher odds of finding a pair in two different isolates than the null expectation. Zero values (white) indicates co-occurrence as expected based on individual frequencies (null expectation). Significance levels: *p ≤ 0.05, **p ≤ 0.01, ***p ≤ 0.001.

## S2 Supplementary Tables

**Table S1:** F-like transfer gene markers, their functions and the number of reference genes used for transfer operon detection. Functional roles have been adapted from [6]. Essential genes used to classifying functional completeness of the F-like transfer operon are indicated with a *.

| F-like tra genes | Function | n reference sequences | Min protein length | Median protein length | Max protein length |
| --- | --- | --- | --- | --- | --- |
| *traM* * | regulation/relaxosome | 27 | 123 | 127 | 168 |
| *traJ* * | regulation/relaxosome | 32 | 67 | 228 | 525 |
| *traY* * | regulation/relaxosome | 22 | 60 | 75 | 133 |
| *traA* * | pilus formation | 25 | 79 | 120 | 132 |
| *traL* * | T4SS | 13 | 98 | 103 | 104 |
| *traE* * | T4SS | 24 | 121 | 188 | 188 |
| *traK* * | T4SS | 24 | 169 | 242 | 475 |
| *traB* * | T4SS | 26 | 422 | 475 | 866 |
| *traP* | nonessential | 27 | 131 | 196 | 208 |
| *trbD* | nonessential | 27 | 63 | 106 | 136 |
| *trbG* | nonessential | 9 | 77 | 83 | 88 |
| *traV* * | T4SS | 27 | 171 | 171 | 204 |
| *traR* * | nonessential | 13 | 57 | 73 | 74 |
| *traC* * | T4SS | 33 | 785 | 876 | 882 |
| *trbI* * | pilus formation | 24 | 101 | 128 | 154 |
| *traW* * | pilus formation | 34 | 195 | 210 | 248 |
| *traU* * | pilus formation | 23 | 322 | 330 | 364 |
| *trbC* | pilus formation | 36 | 200 | 212 | 221 |
| *traN* * | mating pair stability | 36 | 284 | 602 | 651 |
| *trbE* | nonessential | 21 | 71 | 85 | 86 |
| *traF* * | pilus formation | 32 | 247 | 249.5 | 331 |
| *trbA* | nonessential | 17 | 112 | 115 | 115 |
| *artA* | nonessential | 5 | 104 | 104 | 111 |
| *traQ* * | pilus formation | 21 | 78 | 94 | 97 |
| *trbB* * | pilus formation | 24 | 160 | 181 | 203 |
| *trbJ* | nonessential | 24 | 89 | 104 | 147 |
| *trbF* | nonessential | 24 | 37 | 130 | 147 |
| *traH* * | pilus formation | 26 | 426 | 457.5 | 460 |
| *traG* * | T4SS | 35 | 268 | 940 | 984 |
| *traS* | self-transfer prevention | 15 | 60 | 169 | 182 |
| *traT* | self-transfer prevention | 35 | 220 | 243 | 293 |
| *traD* * | T4SS | 47 | 46 | 732 | 1767 |
| *traI* * | regulation/relaxosome | 39 | 130 | 1756 | 1766 |
| *traX* * | pilus formation | 39 | 153 | 248 | 405 |
| *finO* | regulation/relaxosome | 31 | 67 | 186 | 211 |

**Table S2:** Post-hoc comparisons of the association between F-like transfer operon completeness and animal environment using using Pearson’s Chi-squared Test. Positive values indicate greater counts than expected in that category, negative values indicate fewer counts than expected in that category. We report adjusted standardized Pearson residuals and Bonferroni-corrected *p* values.

| Animal Environment | Complete |  | Incomplete |  | Missing |  |
| --- | --- | --- | --- | --- | --- | --- |
|  | Residual | *p* | Residual | *p* | Residual | *p* |
| Bovine | −8.24 | < .001 | −2.37 | 0.162 | 10.53 | < .001 |
| Poultry | 9.29 | < .001 | 1.39 | 1.000 | −10.70 | < .001 |
| Swine | −1.05 | 1.000 | 0.97 | 1.000 | 0.17 | 1.000 |

**Table S3:**
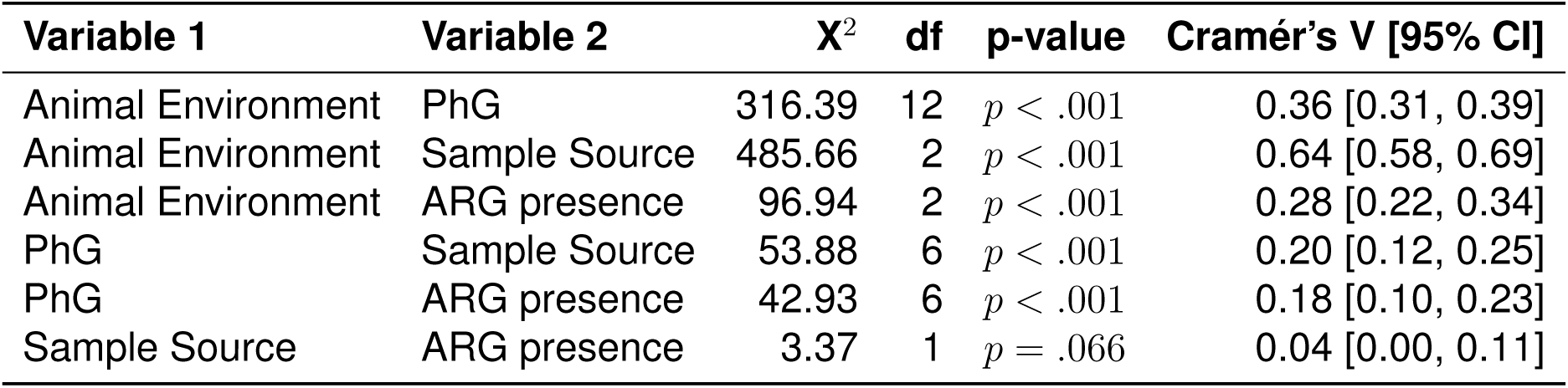
Associations between Animal environment, PhG, Sample Source, and ARG presence tested with Chi-squared Test of Independence. Effect sizes for associations was estimated using Cramér’s V and 95% confidence intervals are reported.

| Variable 1 | Variable 2 | X<sup>2</sup> | df | p-value | Cramér's V [95% CI] |
| --- | --- | --- | --- | --- | --- |
| Animal Environment | PhG | 316.39 | 12 | $p < .001$ | 0.36 [0.31, 0.39] |
| Animal Environment | Sample Source | 485.66 | 2 | $p < .001$ | 0.64 [0.58, 0.69] |
| Animal Environment | ARG presence | 96.94 | 2 | $p < .001$ | 0.28 [0.22, 0.34] |
| PhG | Sample Source | 53.88 | 6 | $p < .001$ | 0.20 [0.12, 0.25] |
| PhG | ARG presence | 42.93 | 6 | $p < .001$ | 0.18 [0.10, 0.23] |
| Sample Source | ARG presence | 3.37 | 1 | $p = .066$ | 0.04 [0.00, 0.11] |

**Table S4:** Selection of main-effects logistic regression model to predict the presence of a complete F-like operon in a genome. All combinations of explanatory variables (Animal Environment, Phylogroup (PhG), Sample Source and ARG presence and Sampling Year) were tested. Models are compared using Akaike Information Criterion (AIC). Lower values of AIC indicate better model fit. The model that includes all Animal Environment, Phylogroup (PhG), Sample Source and ARG presence had the lowest AIC (in bold).

| Model Formula | Residual df | Residual Deviance | AIC |
| --- | --- | --- | --- |
| **Animal Environment + PhG + Sample Source + ARG Presence** | **1189** | **1433** | **1455** |
| Animal Environment + PhG + Sample Source | 1190 | 1439 | 1459 |
| Animal Environment + PhG + Sample Source + Sampling Year | 1189 | 1438 | 1460 |
| Animal Environment + PhG + ARG Presence | 1190 | 1441 | 1461 |
| Animal Environment + PhG + ARG Presence + Sampling Year | 1189 | 1439 | 1461 |
| Animal Environment + PhG | 1191 | 1445 | 1463 |
| Animal Environment + PhG + Sampling Year | 1190 | 1444 | 1464 |
| PhG + Sample Source + ARG Presence | 1191 | 1452 | 1470 |
| PhG + Sample Source + ARG Presence + Sampling Year | 1190 | 1450 | 1470 |
| PhG + Sample Source + Sampling Year | 1191 | 1464 | 1482 |
| PhG + Sample Source | 1192 | 1466 | 1482 |
| Animal Environment + Sample Source + ARG Presence + Sampling Year | 1194 | 1476 | 1488 |
| Animal Environment + Sample Source + ARG Presence | 1195 | 1480 | 1490 |
| Animal Environment + ARG Presence + Sampling Year | 1195 | 1480 | 1490 |
| Animal Environment + ARG Presence | 1196 | 1483 | 1491 |
| Animal Environment + Sample Source + Sampling Year | 1195 | 1484 | 1494 |
| Animal Environment + Sampling Year | 1196 | 1486 | 1494 |
| Animal Environment + Sample Source | 1196 | 1487 | 1495 |
| Animal Environment | 1197 | 1489 | 1495 |
| PhG + ARG Presence | 1192 | 1501 | 1517 |
| PhG + ARG Presence + Sampling Year | 1191 | 1501 | 1519 |
| Phylogenetic Group (PhG) | 1193 | 1510 | 1524 |
| Sample Source + ARG Presence + Sampling Year | 1196 | 1517 | 1525 |
| PhG + Sampling Year | 1192 | 1510 | 1526 |
| Sample Source + ARG Presence | 1197 | 1524 | 1530 |
| Sample Source + Sampling Year | 1197 | 1537 | 1543 |
| Sample Source | 1198 | 1545 | 1549 |
| ARG Presence + Sampling Year | 1197 | 1575 | 1581 |
| ARG Presence | 1198 | 1578 | 1582 |
| Sampling Year | 1198 | 1591 | 1595 |

**Table S5:**
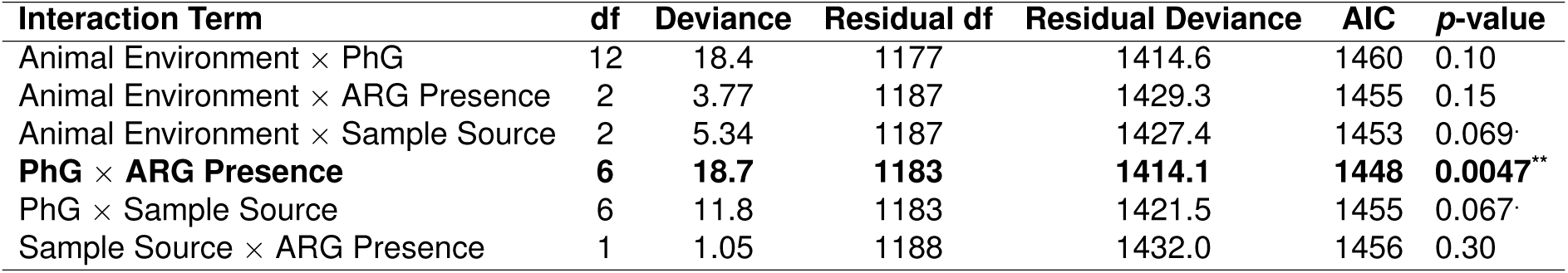
Likelihood ratio tests for interaction terms added to the main effects model containing Animal Environment, PhG, Sample Source and ARG Presence as covariates. The interaction term between PhG and ARG presence significantly improved model fit and was added to the main effects model. This model had the lowest AIC among models tested and was used for subsequent analyses.**\*\*** *p<* 0.01; **\*** *p<* 0.05; . *p<* 0.1.

## S3 Supplementary Text

### S3.1 Genome metadata

The metadata for the genomes were retrieved from [7] where extensive metadata standardization was done to ensure its quality. We describe here briefly the details of the specific metadata variables used in this paper:

- Animal Host: which animal production group the sample is associated with (Poultry, Swine or Bovine)
- Phylogenetic group: the phylogenetic group (PhG) was determined *in silico* using ClermonTyping (version 1.0.0 [8]) and MLST performed with mlst (version 2.16.2 [9, 10]).
- ARG presence: Binary value indicating whether the sample contained antimicrobial resistance genes (ARGs). ARGs were screened using ABRicate (version 0.8.13 [11]) with the ResFinder database (version of 22 April 2019 [12]). If the assembly contained ARG(s), ARG presence was 1.
- Sample Source: Indicates whether the samples were recovered from food products (Food) or from the animals themselves (Livestock). We assume, throughout most of this paper, that the food isolates form part of the broader ecology of the animal environment itself. This is because *E. coli* recovered from food products can be introduced from various sources at multiple stages of the animal production production process [13]. For instance, extra-intestinal pathogenic *Escherichia coli* (ExPEC) commonly found on poultry food samples have been shown to likely stem from the live bird itself rather than from humans [14].
- Sampling Year: Indicates whether the sampling year was before 2017 or 2017 and later (63.2% of isolates).

### S3.2 Reference *tra* gene database construction

We curated a non-redundant reference dataset of amino acid sequence homologues of 35 F-like *tra* genes comprising 917 sequences in total (Supplementary Table S1). The dataset was adapted from Fernandez-Lopez *et al* [15] by removing duplicates, re-annotating the sequences based on updated NCBI protein accession numbers and manually adding missing annotations. The reference *tra* sequences stem from 70 F-like plasmids found in Enterobacteriaceae: *E. coli* (656 *tra* sequences from 48 plasmids), *Salmonella enteric*a (164 *tra* sequences from 14 plasmids), *Salmonella sp.* (24 sequences from 1 plasmid), *Shigella dysenteriae* (2 sequences from 1 plasmid), *Shigella flexneri* (37 sequences from 4 plasmids), *Shigella sonnei*(4 sequences from 1 plasmid) and 33 sequences with an unidentified host.

### S3.3 F-like transfer gene essentiality

In total we detected 35 genes of the F-like transfer operon, of which 26 have a characterized function in conjugation [6, 16, 17]. From this set, 23 can be described as ”essential” for self-transfer (Table S1), i.e. the gene has a demonstrated function in conjugation or its regulation, and mutations in these genes block or reduce plasmid transfer [6, 16, 17]. Entry and surface exclusion genes *traS* and *traT* are not considered essential genes, because they are not involved in the core conjugation machinery and experimental evidence shows that deleting these genes results in increased transfer rates [18, 19]. The fertility inhibition gene *finO* is also not essential for conjugation, as it negatively regulates *tra* gene expression, with higher rates of transfer observed in the classic F plasmid which has a non-functional *finO* gene [6, 20].

### S3.4 Mobility typing scheme from Coluzzi *et al*

We performed mobility typing on each genome using the classification scheme from Coluzzi *et al* [5]. This mobility typing scheme uses the presence of the relaxase and a small number of other conjugation-related genes (including VirB4, T4CP and MPF genes) to classify a plasmid as conjugative (conj; containing the relaxase, VirB4, T4CP and at least 3 MPF proteins), mobilizable (mob; containing the relaxase but missing other components required to be conjugative), non-mobilizable (mobless; lacking relaxases) or partially decayed conjugative (pdconj; a subcategory of MOB plasmids with more than 5 conjugation proteins excluding the relaxase). Again, we used the set of F-like transfer genes we identified in each genome for this classification.

### S3.5 Plasmid reconstruction and clustering

We used MOB-recon from MOB-Suite (v.3.1.9, default parameters, using a set of closed *Enterobacteriaceae* genomes for chromosome depletion) for plasmid reconstruction and clustering. The default mash distance threshold of 0.05 was used to assign plasmid cluster identities. Each genome is represented as a binary vector that describes the presence and absence of these plasmid cluster IDs.

The plasmid clusters specifically associated with the F-like plasmid-associated contigs identified using our transfer gene finding protocol were considered to be F-like plasmid clusters. Only 38.5% of the genomes containing an F-like transfer operon (n=786) could be mapped to an F-like plasmid cluster.

### S3.6 Identifying redundant genomes for sensitivity analyses

Pairwise average nucleotide identity (ANI) was computed using default settings of fas-tANI (v.1.34) [21]. Out of all the genomes sampled at the same time and location, we found 35 clusters of genomes (101 genomes in total) with pairwise ANI ≥99.99%, which is a recommended ANI threshold for strain classification [3]. Out of these clusters, 20 (60 genomes in total) also had similar plasmid content, where pairwise plasmid content similarity was measured using the Jaccard similarity index on the presence/absence vector of plasmid cluster identities (see Section S3.5). Therefore, 40 out of 1200 genomes (0.03% of the total dataset) are presumably redundant. We kept a random representative genome from each of these clusters to form a reduced dataset of 1160 genomes to test the sensitivity of our logistic regression results to redundant genomes.

### S3.7 Co-occurrence of conjugation types

Co-occurrence between two conjugation types in an isolate was assessed using a prevalence-insensitive metric called affinity [22]. For a pair of conjugation types A and B, affinity measures the odds of finding type B in an isolate given that the isolate already has type A. A positive affinity (*>* 0) indicates that types A and B co-occur more frequently than expected under the null distribution, whereas a negative value (*<* 0) indicates that they occur in separate genomes more frequently than expected.

Pairwise affinities were computed for all pairs of conjugation types, which includes 6 pairs of MPF types, 10 pairs of relaxase types and 20 combinations of relaxase and MPF types using the R package CooccurrenceAffinity v.1.0.2 [23]. Since we found only 1 occurrence of MOB V in the entire set of genomes, we excluded this type from the analyses.

## References

1. Carattoli, A. Plasmids and the spread of resistance. International Journal of Medical Microbiology. Special Issue Antibiotic Resistance 303, 298–304 (Aug. 2013).

2. Stoesser, N. et al. Evolutionary History of the Global Emergence of the Escherichia coli Epidemic Clone ST131. mBio 7. Publisher: American Society for Microbiology, 10.1128/mbio.02162-15 (Mar. 2016).

3. Koraimann, G. Spread and Persistence of Virulence and Antibiotic Resistance Genes: A Ride on the F Plasmid Conjugation Module. EcoSal Plus 8. Publisher: American Society for Microbiology (July 2018).

4. Johnson, T. J. & Nolan, L. K. Pathogenomics of the Virulence Plasmids of Escherichia coli. Microbiology and Molecular Biology Reviews 73. Publisher: American Society for Microbiology, 750–774 (Dec. 2009).

5. Yu, M. K., Fogarty, E. C. & Eren, A. M. Diverse plasmid systems and their ecology across human gut metagenomes revealed by PlasX and MobMess. en. Nature Microbiology 9. Publisher: Nature Publishing Group, 830–847 (Mar. 2024).

6. Virolle, C., Goldlust, K., Djermoun, S., Bigot, S. & Lesterlin, C. Plasmid Transfer by Conjugation in Gram-Negative Bacteria: From the Cellular to the Community Level. en. Genes 11. Number: 11 Publisher: Multidisciplinary Digital Publishing Institute, 1239 (Nov. 2020).

7. Firth, N., Ippen-Ihler, K., Skurray, R. A., et al. Structure and function of the F factor and mechanism of conjugation. Escherichia coli and Salmonella: cellular and molecular biology, 2nd ed. ASM Press, Washington, DC, 2377–2401 (1996).

8. Frost, L. S., Ippen-Ihler, K. & Skurray, R. A. Analysis of the sequence and gene products of the transfer region of the F sex factor. Microbiological Reviews 58. Publisher: American Society for Microbiology, 162–210 (June 1994).

9. Huisman, J. S., Bernhard, A. & Igler, C. Should I stay or should I go: transmission trade-offs in phages and plasmids. Trends in Microbiology 33, 484–495 (2025).

10. Bethke, J. H. et al. Vertical and horizontal gene transfer tradeoffs direct plasmid fitness. Molecular Systems Biology 19, e11300 (Feb. 2023).

11. Turner, P. E., Cooper, V. S. & Lenski, R. E. Tradeoff between horizontal and vertical modes of transmission in bacterial plasmids. Evolution 52, 315–329 (Apr. 1998).

12. Millan, A. S. & MacLean, R. C. Fitness Costs of Plasmids: a Limit to Plasmid Transmission. EN. Microbiology Spectrum. Publisher: ASM Press Washington, DC (Sept. 2017).

13. Hall, J. P. J. et al. Plasmid fitness costs are caused by specific genetic conflicts enabling resolution by compensatory mutation. en. PLOS Biology 19. Publisher: Public Library of Science, e3001225 (Oct. 2021).

14. Dahlberg, C. & Chao, L. Amelioration of the Cost of Conjugative Plasmid Carriage in Eschericha coli K12. Genetics 165, 1641–1649 (Dec. 2003).

15. Porse, A., Schønning, K., Munck, C. & Sommer, M. O. Survival and Evolution of a Large Multidrug Resistance Plasmid in New Clinical Bacterial Hosts. Molecular Biology and Evolution 33, 2860–2873 (Nov. 2016).

16. Jalasvuori, M., Friman, V.-P., Nieminen, A., Bamford, J. K. H. & Buckling, A. Bac-teriophage selection against a plasmid-encoded sex apparatus leads to the loss of antibiotic-resistance plasmids. eng. Biology Letters 7, 902–905 (Dec. 2011).

17. Quinones-Olvera, N. et al. Diverse and abundant phages exploit conjugative plasmids. en. Nature Communications 15. Publisher: Nature Publishing Group, 3197 (Apr. 2024).

18. Koraimann, G., Teferle, K., Markolin, G., Woger, W. & Högenauer, G. The FinOP repressor system of plasmid R1: analysis of the antisense RNA control of traJ expression and conjugative DNA transfer. en. Molecular Microbiology 21, 811–821 (1996).

19. Frost, L. S. & Koraimann, G. Regulation of bacterial conjugation: balancing opportunity with adversity. Future Microbiology 5. Publisher: Future Medicine, 1057–1071 (July 2010).

20. Achtman, M., Kennedy, N. & Skurray, R. Cell–cell interactions in conjugating Escherichia coli: role of traT protein in surface exclusion. en. Proceedings of the National Academy of Sciences 74, 5104–5108 (Nov. 1977).

21. Rivard, N. et al. Surface exclusion of IncC conjugative plasmids and their relatives. en. PLOS Genetics 20. Publisher: Public Library of Science, e1011442 (Oct. 2024).

22. Coluzzi, C., Garcillán-Barcia, M. P., de la Cruz, F. & Rocha, E. P. Evolution of Plasmid Mobility: Origin and Fate of Conjugative and Nonconjugative Plasmids. Molecular Biology and Evolution 39, msac115 (June 2022).

23. Hanke, D. M., Wang, Y. & Dagan, T. Pseudogenes in plasmid genomes reveal past transitions in plasmid mobility en. Pages: 2023.11.08.566193 Section: New Results. Apr. 2024.

24. Fernandez-Lopez, R., de Toro, M., Moncalian, G., Garcillan-Barcia, M. P. & de la Cruz, F. Comparative Genomics of the Conjugation Region of F-like Plasmids: Five Shades of F. Frontiers in Molecular Biosciences 3 (2016).

25. Zhang, F. et al. Comparative genomics reveals new insights into the evolution of the IncA and IncC family of plasmids. English. Frontiers in Microbiology 13. Publisher: Frontiers (Nov. 2022).

26. Benz, F. et al. Plasmid- and strain-specific factors drive variation in ESBL-plasmid spread in vitro and in vivo. en. The ISME Journal 15. Number: 3 Publisher: Nature Publishing Group, 862–878 (Mar. 2021).

27. Douarre, P.-E., Mallet, L., Radomski, N., Felten, A. & Mistou, M.-Y. Analysis of COMPASS, a New Comprehensive Plasmid Database Revealed Prevalence of Multireplicon and Extensive Diversity of IncF Plasmids. Frontiers in Microbiology 11 (2020).

28. Ares-Arroyo, M., Coluzzi, C. & Rocha, E. P. C. Origins of transfer establish networks of functional dependencies for plasmid transfer by conjugation. Nucleic Acids Research 51, 3001–3016 (Apr. 2023).

29. Domingues, C. P. F., Rebelo, J. S., Dionisio, F. & Nogueira, T. Clinical and Environmental Plasmids: Antibiotic Resistance, Virulence, Mobility, and ESKAPEE Pathogens. Antibiotics 15, 29 (Dec. 2025).

30. Matlock, W. et al. Genomic network analysis of environmental and livestock F-type plasmid populations. en. The ISME Journal 15. Number: 8 Publisher: Nature Publishing Group, 2322–2335 (Aug. 2021).

31. Shaw, L. P. et al. Niche and local geography shape the pangenome of wastewater-and livestock-associated Enterobacteriaceae. Science Advances 7. Publisher: American Association for the Advancement of Science, eabe3868 (Apr. 2021).

32. Stephens, C. et al. F Plasmids Are the Major Carriers of Antibiotic Resistance Genes in Human-Associated Commensal Escherichia coli. EN. mSphere. Publisher: American Society for Microbiology1752 N St., N.W., Washington, DC (Aug. 2020).

33. Reid, C. J., Cummins, M. L. & Djordjevic, S. P. Major F plasmid clusters are linked with ColV and pUTI89-like marker genes in bloodstream isolates of Escherichia coli. BMC Genomics 26, 57 (Jan. 2025).

34. Garcillán-Barcia, M. P., Francia, M. V. & de La Cruz, F. The diversity of conjugative relaxases and its application in plasmid classification. FEMS Microbiology Reviews 33, 657–687 (May 2009).

35. Finks, S. S. & Martiny, J. B. H. Plasmid-Encoded Traits Vary across Environments. mBio 14. Publisher: American Society for Microbiology, e03191–22 (Jan. 2023).

36. Smillie, C., Garcillán-Barcia, M. P., Francia, M. V., Rocha, E. P. C. & de la Cruz, F. Mobility of Plasmids. Microbiology and Molecular Biology Reviews 74. Publisher: American Society for Microbiology, 434–452 (Sept. 2010).

37. Garcillán-Barcia, M. P., de la Cruz, F. & Rocha, E. P. C. The extended mobility of plasmids. Nucleic Acids Research 53, gkaf652 (Aug. 2025).

38. Pires, J., Huisman, J. S., Bonhoeffer, S. & Van Boeckel, T. P. Increase in antimicrobial resistance in Escherichia coli in food animals between 1980 and 2018 assessed using genomes from public databases. Journal of Antimicrobial Chemotherapy 77, 646–655 (Mar. 2022).

39. Achtman, M., Zhou, Z., Charlesworth, J. & Baxter, L. EnteroBase: hierarchical clustering of 100 000s of bacterial genomes into species/subspecies and populations. Philosophical Transactions of the Royal Society B: Biological Sciences 377. Publisher: Royal Society, 20210240 (Aug. 2022).

40. Zhou, Z., Alikhan, N.-F., Mohamed, K., Fan, Y. & Achtman, M. The EnteroBase user’s guide, with case studies on Salmonella transmissions, Yersinia pestis phylogeny, and Escherichia core genomic diversity. Genome Research 30. eprint: http://genome.cshlp.org/content/genome/30/1/138.full.pdf, 138–152 (2020).

41. Chklovski, A., Parks, D. H., Woodcroft, B. J. & Tyson, G. W. CheckM2: a rapid, scalable and accurate tool for assessing microbial genome quality using machine learning. en. Nature Methods 20, 1203–1212 (Aug. 2023).

42. Camacho, C. et al. BLAST+: architecture and applications. eng. BMC bioinformatics 10, 421 (Dec. 2009).

43. Seddon, C. et al. Cryo-EM structure and evolutionary history of the conjugation surface exclusion protein TraT. en. Nature Communications 16. Publisher: Nature Publishing Group, 659 (Jan. 2025).

44. Schwengers, O., et al. Bakta: rapid and standardized annotation of bacterial genomes via alignment-free sequence identification. Microbial Genomics 7. Publisher: Microbiology Society, 000685 (2021).

45. Garcillán-Barcia, M. P. & de la Cruz, F. Why is entry exclusion an essential feature of conjugative plasmids? en. Plasmid 60, 1–18 (July 2008).

46. Carattoli, A. & Hasman, H. en. in Horizontal Gene Transfer: Methods and Protocols (ed de la Cruz, F.) 285–294 (Springer US, New York, NY, 2020).

47. Lenth, R. V. & Piaskowski, J. emmeans: Estimated Marginal Means, aka Least-Squares Means en. Institution: Comprehensive R Archive Network Pages: 2.0.1. Oct. 2017.

48. Camargo, A. P. et al. Identification of mobile genetic elements with geNomad. en. Nature Biotechnology 42. Publisher: Nature Publishing Group, 1303–1312 (Aug. 2024).

49. Rodriguez-R, L. M. et al. An ANI gap within bacterial species that advances the definitions of intra-species units. mBio 15, e02696–23 (Dec. 2023).

50. Viver, T. et al. Towards estimating the number of strains that make up a natural bacterial population. en. Nature Communications 15, 544 (Jan. 2024).

51. Touchon, M. et al. Phylogenetic background and habitat drive the genetic diversification of Escherichia coli. en. PLOS Genetics 16. Publisher: Public Library of Science, e1008866 (June 2020).

52. Orlek, A. et al. Plasmid Classification in an Era of Whole-Genome Sequencing: Application in Studies of Antibiotic Resistance Epidemiology. Frontiers in Microbiology 8 (2017).

53. Arredondo-Alonso, S., Willems, R. J., van Schaik, W. & Schürch, A. C. On the (im)possibility of reconstructing plasmids from whole-genome short-read sequencing data. Microbial Genomics 3. Publisher: Microbiology Society, e000128 (2017).

54. San Millan, A., Heilbron, K. & MacLean, R. C. Positive epistasis between co-infecting plasmids promotes plasmid survival in bacterial populations. The ISME Journal 8, 601–612 (Mar. 2014).

55. Mainali, K. P., Slud, E., Singer, M. C. & Fagan, W. F. A better index for analysis of co-occurrence and similarity. Science Advances 8. Publisher: American Association for the Advancement of Science, eabj9204 (Jan. 2022).

56. Moser, K. A. et al. The Role of Mobile Genetic Elements in the Spread of Antimicrobial-Resistant Escherichia coli From Chickens to Humans in Small-Scale Production Poultry Operations in Rural Ecuador. American Journal of Epidemiology 187, 558–567 (Mar. 2018).

57. Rozwandowicz, M. et al. Successful Host Adaptation of IncK2 Plasmids. English. Frontiers in Microbiology 10 (Oct. 2019).

58. Marmion, M., Ferone, M. T., Whyte, P. & Scannell, A. G. M. The changing microbiome of poultry meat; from farm to fridge. Food Microbiology 99, 103823 (Oct. 2021).

59. Tardiolo, G. et al. Gut Microbiota of Ruminants and Monogastric Livestock: An Overview. Animals : an Open Access Journal from MDPI 15, 758 (Mar. 2025).

60. Klimke, W. A. & Frost, L. S. Genetic Analysis of the Role of the Transfer Gene, *traN*, of the F and R100-1 Plasmids in Mating Pair Stabilization during Conjugation. en. Journal of Bacteriology 180, 4036–4043 (Aug. 1998).

61. Lu, J. et al. Structural basis of specific TraD–TraM recognition during F plasmid-mediated bacterial conjugation. en. Molecular Microbiology 70, 89–99 (2008).

62. Maneewannakul, K. et al. Construction of derivatives of the F plasmid pOX–tra715: characterization of traY and traD mutants that can be complemented in trans. en. Molecular Microbiology 22. eprint: https://onlinelibrary.wiley.com/doi/pdf/10.1046/j.1365-2958.1996.00087.x, 197–205 (1996).

63. Kishida, K. et al. Chimeric systems composed of swapped Tra subunits between distantly-related F plasmids reveal striking plasticity among type IV secretion machines. en. PLOS Genetics 20, e1011088 (Mar. 2024).

## References

1. Chklovski, A., Parks, D. H., Woodcroft, B. J. & Tyson, G. W. CheckM2: a rapid, scalable and accurate tool for assessing microbial genome quality using machine learning. en. Nature Methods 20, 1203–1212 (Aug. 2023).

2. Parks, D. H., Imelfort, M., Skennerton, C. T., Hugenholtz, P. & Tyson, G. W. CheckM: assessing the quality of microbial genomes recovered from isolates, single cells, and metagenomes. Genome Research 25, 1043–1055 (July 2015).

3. Viver, T. et al. Towards estimating the number of strains that make up a natural bacterial population. en. Nature Communications 15, 544 (Jan. 2024).

4. Rodriguez-R, L. M. et al. An ANI gap within bacterial species that advances the definitions of intra-species units. mBio 15, e02696–23 (Dec. 2023).

5. Coluzzi, C., Garcillán-Barcia, M. P., de la Cruz, F. & Rocha, E. P. Evolution of Plasmid Mobility: Origin and Fate of Conjugative and Nonconjugative Plasmids. Molecular Biology and Evolution 39, msac115 (June 2022).

6. Koraimann, G. Spread and Persistence of Virulence and Antibiotic Resistance Genes: A Ride on the F Plasmid Conjugation Module. EcoSal Plus 8. Publisher: American Society for Microbiology (July 2018).

7. Pires, J., Huisman, J. S., Bonhoeffer, S. & Van Boeckel, T. P. Increase in antimicrobial resistance in Escherichia coli in food animals between 1980 and 2018 assessed using genomes from public databases. Journal of Antimicrobial Chemotherapy 77, 646–655 (Mar. 2022).

8. Beghain, J., Bridier-Nahmias, A., Nagard, H. L., Denamur, E. & Clermont, O. ClermonTyping: an easy-to-use and accurate in silico method for Escherichia genus strain phylotyping. en. Microbial Genomics 4, e000192 (July 2018).

9. Jolley, K. A., Bray, J. E. & Maiden, M. C. J. Open-access bacterial population genomics: BIGSdb software, the PubMLST.org website and their applications. eng. Wellcome Open Research 3, 124 (2018).

10. Seemann, T. tseemann/mlst original-date: 2014-05-03T09:12:11Z. July 2026.

11. Seemann, T. tseemann/abricate original-date: 2014-07-17T00:59:35Z. July 2026.

12. Bortolaia, V. et al. ResFinder 4.0 for predictions of phenotypes from genotypes. eng. The Journal of Antimicrobial Chemotherapy 75, 3491–3500 (Dec. 2020).

13. Risalvato, J., Sewid, A. H., Eda, S., Gerhold, R. W. & Wu, J. J. Strategic Detection of Escherichia coli in the Poultry Industry: Food Safety Challenges, One Health Approaches, and Advances in Biosensor Technologies. en. Biosensors 15, 419 (July 2025).

14. Bergeron, C. R. et al. Chicken as Reservoir for Extraintestinal Pathogenic Escherichia coli in Humans, Canada - Volume 18, Number 3—March 2012 - Emerging Infectious Diseases journal - CDC. en-us.

15. Fernandez-Lopez, R., de Toro, M., Moncalian, G., Garcillan-Barcia, M. P. & de la Cruz, F. Comparative Genomics of the Conjugation Region of F-like Plasmids: Five Shades of F. Frontiers in Molecular Biosciences 3 (2016).

16. Frost, L. S., Ippen-Ihler, K. & Skurray, R. A. Analysis of the sequence and gene products of the transfer region of the F sex factor. Microbiological Reviews 58. Publisher: American Society for Microbiology, 162–210 (June 1994).

17. Firth, N., Ippen-Ihler, K., Skurray, R. A., et al. Structure and function of the F factor and mechanism of conjugation. Escherichia coli and Salmonella: cellular and molecular biology, 2nd ed. ASM Press, Washington, DC, 2377–2401 (1996).

18. Achtman, M., Kennedy, N. & Skurray, R. Cell–cell interactions in conjugating Escherichia coli: role of traT protein in surface exclusion. en. Proceedings of the National Academy of Sciences 74, 5104–5108 (Nov. 1977).

19. Couturier, A., Fraikin, N. & Lesterlin, C. Exclusion systems preserve host cell homeostasis and fitness, ensuring successful dissemination of conjugative plasmids and associated resistance genes. Nucleic Acids Research 53, gkaf898 (Sept. 2025).

20. Zatyka, M. & Thomas, C. M. Control of genes for conjugative transfer of plasmids and other mobile elements. FEMS Microbiology Reviews 21, 291–319 (Feb. 1998).

21. Jain, C., Rodriguez-R, L. M., Phillippy, A. M., Konstantinidis, K. T. & Aluru, S. High throughput ANI analysis of 90K prokaryotic genomes reveals clear species boundaries. en. Nature Communications 9, 5114 (Nov. 2018).

22. Mainali, K. P., Slud, E., Singer, M. C. & Fagan, W. F. A better index for analysis of co-occurrence and similarity. Science Advances 8. Publisher: American Association for the Advancement of Science, eabj9204 (Jan. 2022).

23. Mainali, K. P. & Slud, E. CooccurrenceAffinity: An R package for computing a novel metric of affinity in co-occurrence data that corrects for pervasive errors in traditional indices. en. PLOS ONE 20. Publisher: Public Library of Science, e0316650 (Jan. 2025).

